# Tumor Suppressor p53 Restrains Cancer Cell Dissemination by Modulating Mitochondrial Dynamics

**DOI:** 10.1101/2021.09.09.459707

**Authors:** Thi Tuyet Trinh Phan, Yu-Chun Lin, Yu-Ting Chou, Chien-Wei Wu, Lih-Yuan Lin

**Affiliations:** Institute of Molecular and Cellular Biology, College of Life Science, National Tsing Hua University, Hsinchu 300044, Taiwan R.O.C; Institute of Molecular Medicine, College of Life Science, National Tsing Hua University, Hsinchu 300044, Taiwan R.O.C; Institute of Biotechnology, College of Life Science, National Tsing Hua University, Hsinchu 300044, Taiwan R.O.C

**Keywords:** p53, migration and invasion, mitochondrial dynamics, ERK1/2, MMP9, EMT

## Abstract

Tumor suppressor p53 plays a central role in preventing tumorigenesis. Here, we unravel how p53 modulates mitochondrial dynamics to restrain the metastatic properties of cancer cells. p53 inhibits the mammalian target of rapamycin complex 1 (mTORC1) signaling to attenuate the protein level of mitochondrial fission process 1 (MTFP1), which fosters the pro-fission dynamin-related protein 1 (Drp1) phosphorylation. This regulatory mechanism allows p53 to restrict cell migration and invasion governed by Drp1-mediated mitochondrial fission. Downregulating p53 expression or elevating the molecular signature of mitochondrial fission correlates with aggressive tumor phenotypes and poor prognosis in cancer patients. Upon p53 loss, exaggerated mitochondrial fragmentation stimulates the activation of the extracellular signal-regulated kinase 1/2 (ERK1/2) signaling resulting in epithelial-to-mesenchymal transition (EMT)-like changes in cell morphology, accompanied by accelerated matrix metalloproteinase-9 (MMP9) expression and invasive cell migration. Notably, blocking the activation of mTORC1/MTFP1/Drp1/ERK1/2 axis completely abolishes the p53 deficiency-driven cellular morphological switch, MMP9 expression, and cancer cell dissemination. Our findings unveil a hitherto unrecognized mitochondria-dependent molecular mechanism underlying the metastatic phenotypes of p53-compromised cancers.

## Introduction

Metastasis is the dissemination of cancer cells from an original primary site to distant organs or tissues resulting in the formation of new tumors within the body [1]. The metastatic cascade of solid tumors can be dissected into five sequential steps: the process initiates with local invasion across basement membrane, follows by intravasation into the blood and lymphatic vessels, continues with survival in the circulatory system, advances with extravasation from the bloodstream to distant sites, and finalizes with colonization at secondary metastatic sites [2]. Cell migration and invasion, which are governed by cell motility, are recognized as key parameters crucial for the initiation and progression of the metastatic cascade [3]. Despite its importance in determining the clinical prognosis of cancer patients, little is known about the rate-limiting steps governing and/or promoting tumor cell motility and invasiveness.

Mitochondria are double membraned subcellular organelles that perform diverse functions in eukaryotic cells. Besides generating metabolic energy to power cellular functions, mitochondria participate in a multitude of vital processes regulating cellular redox, calcium homeostasis, aging, and cell death [4, 5]. Recently, mitochondrial dynamics has been implicated in controlling the metastatic dissemination of cancer cells [6–11]. Enforcing mitochondrial fission or inhibiting mitochondrial fusion supports cell migration, invasion, and metastasis in hepatocellular carcinoma, glioma, pancreatic, breast, and bladder cancers. Paradoxically, increased mitochondrial fission attenuates metastasis in triple-negative breast cancer [12]. Thus, whether mitochondrial fission or fusion advances the metastatic potential of cancer cells may be context-dependent and requires further investigation.

Mitochondrial fission and fusion processes are controlled by a conserved superfamily of dynamin-like guanosine triphosphatases (GTPases) [13]. Mitochondrial fusion is mediated by Mitofusin 1 (Mfn1), Mitofusin 2 (Mfn2), and Optic Atrophy 1 (Opa1). Mitochondria fission is controlled by dynamin-related protein 1 (Drp1) and its accessory receptors mitochondrial fission 1 (Fis1), mitochondrial fission factor (Mff), and mitochondrial dynamics proteins of 49 and 51 kDa (MiD49/MIEF2 and MiD51/MIEF1) [14]. Phosphorylation of Drp1 at serine 616 (S616) and serine 637 (S637) is best known for regulating Drp1 activity and mitochondrial fission. Phosphorylation of S637 on Drp1 inhibits its GTPase activity, promoting the elongation of the mitochondrial network, whereas phosphorylation of S616 on Drp1 facilitates the translocation of cytosolic Drp1 to the outer mitochondrial membrane and induces mitochondrial fragmentation [15]. Because mitochondrial dynamics is tightly associated with the metastatic abilities of cancer cells, identification of additional regulators modulating mitochondrial dynamics could benefit the discovery of novel therapeutic approaches to target malignant tumors.

Mammalian target of rapamycin (mTOR) is a serine/threonine-protein kinase that functions as the catalytic subunit of two distinct multiprotein complexes, named mTOR complex 1 (mTORC1) and mTOR complex 2 (mTORC2) [16]. It has been established that mTORC1 is an important regulator of mitochondrial dynamics. mTORC1 phosphorylates 4EBPs (the translation initiation factor 4E (eIF4E)-binding proteins) and prevents it from binding eIF4E. The eIF4E can then initiate the translation of the mitochondrial fission process 1 (MTFP1). MTFP1 is a transmembrane protein locates in the mitochondrial inner membrane and facilitates Drp1-driven mitochondrial fission [17, 18]. Studies have suggested mTOR activation contributes to elevated cancer migration, invasion, and metastasis [19, 20], while mTOR inhibition results in mitochondrial elongation and branching [17, 21] and the morphology can be completely reversed by overexpressing MTFP1 [17]. However, the links among mTOR, metastasis, and mitochondrial dynamics have not been examined.

The tumor suppressor p53, encoded by the tumor protein p53 (*TP53*) gene, is a master regulator of multiple cell fate-determining genes and prevents the oncogenic activation of the mTOR signaling pathway [22–25]. Loss of p53 activity is a hallmark of most human tumors, affecting more than one-half of cancer cases [26]. Accumulating data suggests an unconventional role of p53 in controlling cancer cell invasiveness [27]. p53 also impacts mitochondrial integrity in response to various stresses by either regulating proteins involved in mitochondrial quality control and metabolism or maintaining the mitochondrial genomic integrity [28–31]. Nonetheless, how p53 modulates the morphological dynamics of mitochondria remains poorly understood. Moreover, whether mitochondrial dynamics is involved in p53-dependent regulation of cell motility and invasion has not been addressed.

In this study, we delineate a p53-regulated circuitry that restrains the metastatic dissemination of cancer cells and contributes to cancer phenotypes and patient prognosis. We show that p53 alleviates the dynamin-related protein 1 (Drp1)-driven mitochondrial fission by inhibiting the mTORC1-mediated MTFP1 protein expression. p53 deficiency-exaggerated mitochondrial fragmentation activates the extracellular signal-regulated kinase 1/2 (ERK1/2) signaling leading to remarkable changes in cell morphology and robust increases in the matrix metalloproteinase 9 (MMP9) expression and invasive cell migration. Hence, mitochondrial fission represents a driving force for signal transduction that directs cancer cell migration and invasion when wild-type (WT) p53 functions are impaired.

## Materials and Methods

### Cell lines and cell culture conditions

A549 and H1299 cells were maintained in Roswell Park Memorial Institute (RPMI) 1640 medium supplemented with 10% heat-inactivated fetal bovine serum (FBS), 0.22% sodium bicarbonate, 2 mM L-glutamine (L-Gln), and 100 units/ml penicillin/streptomycin (P/S). MCF-7 cells were maintained in Dulbecco’s modified Eagle’s medium (DMEM) containing 10% FBS, 0.37% sodium bicarbonate, 2 mM L-Gln, and 100 units/ml P/S. All cells were cultured at 37 °C in a humidified incubator supplemented with 5% CO_2_.

### Reagents and treatments

Reagents for cell cultures were purchased from Invitrogen Gibco (Grand Island, NY, USA). Other chemicals in this study were purchased from Sigma-Aldrich (St. Louis, MO, USA) unless specified. Sodium arsenite was obtained from Merck (Darmstadt, Germany). MTT (3-(4,5-dimethylthiazol-2-yl-2, 5-diphenyl tetrazolium bromide)) was purchased from Alfa Aesar (Thermo Fisher Scientific, Leicestershire, UK). MitoTracker Green FM, MitoSOX Red, JC-1 dye, TRIzol reagent, reagents for reverse transcription and transfection, and siRNAs were purchased from Invitrogen (Carlsbad, CA, USA). Matrigel basement membrane matrix was purchased from Corning (Tewksbury, MA, USA). Primers used in qRT-PCR were purchased from Integrated DNA Technologies (Coralville, IA, USA). PD98059 was purchased from Enzo Life Sciences (Farmingdale, NY, USA).

Sodium arsenite (SA) (Merck) was dissolved in deionized water and added to culture medium at indicated concentrations (10, 20, 40, and 80 µM) for 24 h. Pifithrin-α (PFT-α) (Sigma-Aldrich) and PD98059 (Enzo Life Sciences) were dissolved in dimethyl sulfoxide (DMSO) (Sigma-Aldrich). Cells were pretreated with PFT-α or PD98059 at 20 µM for 3 h or 30 µM for 2 h, respectively, prior to treatment with 20 µM SA for 24 h.

### siRNA and plasmid transfection

Small interference (si)RNA-mediated gene knockdown experiments were carried out with cells transfected with 10 nM siRNA (Invitrogen) for 3 days using Lipofectamine RNAimax (Invitrogen) following the manufacturer’s reverse transfection instructions. A Stealth RNAi siRNA Negative Control (Invitrogen) was used as control. See Table S1 for information about siRNA target sequences.

The pcDNA3 p53 WT plasmid was constructed by inserting a WT p53 gene (393 amino acids) into a pcDNA3 plasmid. Transient expression of WT p53 in H1299 cells was carried out by transfecting cells with the pcDNA3 p53 WT plasmids for 2 days using Lipofectamine 2000 reagent (Invitrogen) according to the manufacturer’s protocol. An empty pcDNA3 plasmid was used as control.

### Cell lysis, immunoblotting, and antibodies

Cells were harvested by trypsinization and centrifugation at 700 × *g* for 5 min at 4 °C and lysed in ice-cold radioimmunoprecipitation assay (RIPA) lysis buffer (50 mM Tris-HCl (pH8.0), 150 mM NaCl, 5 mM EDTA (pH 8.0), 1% NP-40, 0.1% SDS, and 0.5% sodium deoxycholate) supplemented with complete protease and phosphatase inhibitor cocktails (Fivephoton Biochemicals, San Diego, CA, USA). Cell suspension was subsequently incubated on ice for 10 min and then vortexed vigorously for 5 sec. The incubation and vortexing steps were repeated four times, and cell lysates were separated from debris by centrifugation at 12 000 × *g* for 20 min at 4 °C. After centrifugation, the supernatants were transferred to new tubes and protein concentrations were determined using the Bio-Rad protein assay (Bio-Rad, Hercules, CA, USA). Proteins in cell lysates were separated on SDS-PAGE and then transferred electrophoretically onto PVDF membranes (GE Healthcare, Milwaukee, WI, USA) using a transfer cell (Bio-Rad, Hercules, CA, USA). Membranes were subsequently pre-hybridized in TBST buffer (150 mM NaCl, 10 mM Tris-HCl (pH 8.0), 0.1% Tween-20) with 5% skim milk for 1 h before incubating overnight with appropriate primary antibodies diluted in TBST buffer containing 5% bovine serum albumin (BSA). Antibodies against p53 (GTX70214) and GAPDH (GTX100118) were purchased from GeneTex (Hsinchu, Taiwan). Antibodies against Drp1 (8570), phospho-Drp1 (Ser616) (3455), phospho-Drp1 (Ser637) (4867), Mfn2 (11925), 4EBP1 (9644), phospho-4EBP1 (Ser65) (9456), p70 S6K (2708), phospho-p70 S6K (Thr389) (9234), mTOR (2972), phospho-mTOR (Ser2448) (2971), ERK1/2 (9102), and phospho-ERK1/2 (Thr202/Tyr204) (4370) were purchase from Cell Signaling Technology (Beverly, MA, USA). Antibodies against MTFP1 (ab198217) were from Abcam (Cambridge, UK). Antibodies against Caspase-2 (MAB3507) and MDM2 (MABE340) were from EMD Millipore Corporation (Burlington, NC, USA). All antibodies were used at a 1:1 000 dilution except anti-GAPDH (1:10 000 dilution) and anti-MTFP1 (1:500 dilution).

After primary antibody incubation, membranes were washed three times for 15 min each with TBST buffer, then incubated for 1 h with the respective horseradish peroxidase (HRP)-conjugated secondary antibodies diluted to 5 000 folds in TBST buffer containing 5% skim milk. HRP-conjugated anti-rabbit IgG (NA934V) and anti-mouse IgG (NA931V) were from Amersham (GE Healthcare, Buckinghamshire, UK). HRP-conjugated rabbit anti-rat IgG (ab6734) was from Abcam (Cambridge, UK). Membranes were then washed three times for 15 min each with TBST buffer and detected by chemiluminescence using CyECL Western Blotting Substrate H (Cyrusbioscience, MDBio, Taipei, Taiwan).

Visualization was processed with an ImageQuant LAS 4000 mini biomolecular imager (GE Healthcare), and the intensities of bands were quantified with the UN-SCAN-IT gel analysis software (version 6.1) (Silk Scientific, Orem, UT, USA). Signal intensities of total proteins were normalized to GAPDH. Signal intensities of phosphorylated proteins were calculated by dividing the GAPDH-normalized signal intensity for each phosphorylated protein by the GAPDH-normalized intensity of the corresponding total protein.

### Live single-cell tracking

The single-cell motility assay was performed as described previously [32]. Briefly, 1×10^4^ cells were plated in 6-well cell culture plates (Corning) and incubated in 5% CO_2_ at 37 °C. Time-lapse microscopy was performed using an LS620 Microscope (Lumascope, San Diego, CA, USA). Live-cell images were taken automatically every 10 min over a 24 h period. Single-cell migration distance and trajectory were analyzed at two different X-Y positions using the “Manual Tracking” plugin of the ImageJ software (NIH, Bethesda, MD, USA).

### *In vitro* wound-healing assay

The scratch wound-healing assay was performed as previously described [33]. Briefly, cells were grown to confluence in 6-well cell culture plates (Corning). A linear wound was created by scratching the cell monolayer with a sterile p200 pipette tip. Cells were washed several times before incubating in an FBS-free medium. To monitor the migration of cells back into the wound area, cells were imaged at 0 h and 24 h after scratching using a Dino-Eye Microscope Eyepiece Camera (AnMo Electronics Corporation, New Taipei City, Taiwan) connected to a Nikon TMS-F Inverted Phase Contrast Microscope (Nikon, Tokyo, Japan). The area of the wound was quantified by the “Wound Healing Tool” plugin of the ImageJ software (NIH, Bethesda, MD, USA). The relative migration into the wound was calculated by normalizing the measured wound closure area to the area of the initial wound at the 0 h time point.

### Transwell cell migration and invasion assays

Transwell cell migration assays were performed using 24-well cell culture inserts with an 8.0-μm pore size transparent polyethylene terephthalate (PET) membrane (Corning) according to the manufacturer’s recommendations. Briefly, 2.5 × 10^5^ cells in 200 μl of serum-free medium were loaded into the upper chamber of each insert. The bottom wells were filled with 750 μl of complete medium containing 10% FBS. The cells were cultured in a humidified incubator at 37 °C supplemented with 5% CO_2_. After 16 h, nonmigratory cells on the upper surface of the inserts were carefully removed with cotton swabs. Migrated cells attached to the lower surface of the inserts were fixed with 4% paraformaldehyde (PFA) (Electron Microscopy Sciences, Hatfield, PA, USA) for 2 min, permeabilized with 100% methanol for 20 min, then stained with 0.05% crystal violet (Sigma-Aldrich) for 15 min at room temperature. Migrated cells were observed and imaged using a Dino-Eye Microscope Eyepiece Camera (AnMo Electronics Corporation, New Taipei City, Taiwan) connected to a Nikon TMS-F Inverted Phase Contrast Microscope (Nikon, Tokyo, Japan). The bound crystal violet was eluted with 33% acetic acid (Mallinckrodt Chemicals, Phillipsburg, NJ, USA) and quantified by measuring the absorbance at 595 nm with a microplate reader (Bio-Rad, Hercules, CA, USA). The relative ability of migration was calculated as folds changed in absorbance of the indicated sample in relation to the control. In the invasion assay, 5 × 10^5^ cells were plated onto the Matrigel-coated inserts and the same procedure as described above was followed.

### Live-cell fluorescence microscopy and quantification of mitochondrial morphology

1×10^4^ A549 cells were grown in poly(D-lysine)-coated borosilicate glass Lab-Tek 8-well chambers (Thermo Scientific) and stained with MitoTracker Green FM (Invitrogen) (50 nM) for 30 min. After staining, cells were washed three times in prewarmed PBS and replaced with fresh prewarmed medium. Live-cell fluorescence images were acquired on a Nikon Eclipse Ti inverted microscope with a 60X oil objective lens (Nikon) and DS-Qi2 CMOS camera (Nikon) using Nikon element AR software (Nikon). Fluorescence images with 15 stacks of 0.3 µm each were deconvoluted using Huygens Essential Software (Scientific Volume Imaging, Hilversum, North Holland, Netherlands). The maximum intensity projections of images were generated by Nikon element AR software (Nikon). Images were mainly processed and analyzed using Nikon element AR software (Nikon) and mitochondrial morphology was classified as elongated, intermediate, or fragmented. Elongated mitochondria were those that have long tubulating or spreading reticular networks. Cells displaying mitochondria that are small spherical, ovoid, or short rod-shaped were presented as fragmented. Cells containing a mixture of small spherical and shorter tubular mitochondria were classified as intermediate.

### RNA isolation and quantitative real-time PCR (qRT-PCR)

Total RNA was extracted using TRIzol reagent (Invitrogen) following the procedures provided by the manufacturer. The extracted RNA was reverse-transcribed with a RevertAid First Strand cDNA Synthesis Kit (Invitrogen). The resulting complementary (c)DNA was used for quantitative real-time PCR (qRT-PCR) using SYBR Green PCR Master Mix (Applied Biosystems, Foster City, CA, USA) on a StepOnePlus Real-Time PCR system (Applied Biosystems). Data were acquired and analyzed using StepOne Software v2.3 (Thermo Fisher Scientific). The expression of glyceraldehyde 3-phosphate dehydrogenase (GAPDH) was determined in each sample and used as reference gene. Expression of target genes was compared on the basis of equivalent GAPDH transcripts using the 2^-ΔΔCt^ method. See Table S2 for information about primer sequences.

### Software and statistical analysis

All analyzes of clinical data carried out in this paper are based upon data generated by The Cancer Genome Atlas (TCGA) Research Network (https://www.cancer.gov/tcga) except the association between the presence of TP53 mutations versus the probability of distant metastasis (MSK-IMPACT cohort) (Fig. 1B). See Supplementary Materials and Methods for details. Data were presented as means ± SD or SEM of at least three independent experiments. All graphing and statistical analyses were performed using GraphPad Prism 7 software. Details of sample size (n), statistical test, and *p-value* applied for each experiment were indicated in the figure legends. Values with *p* < 0.05 and designated with * are considered statistically significant. All composite figures were assembled in Adobe Illustrator.

**Fig. 1.**
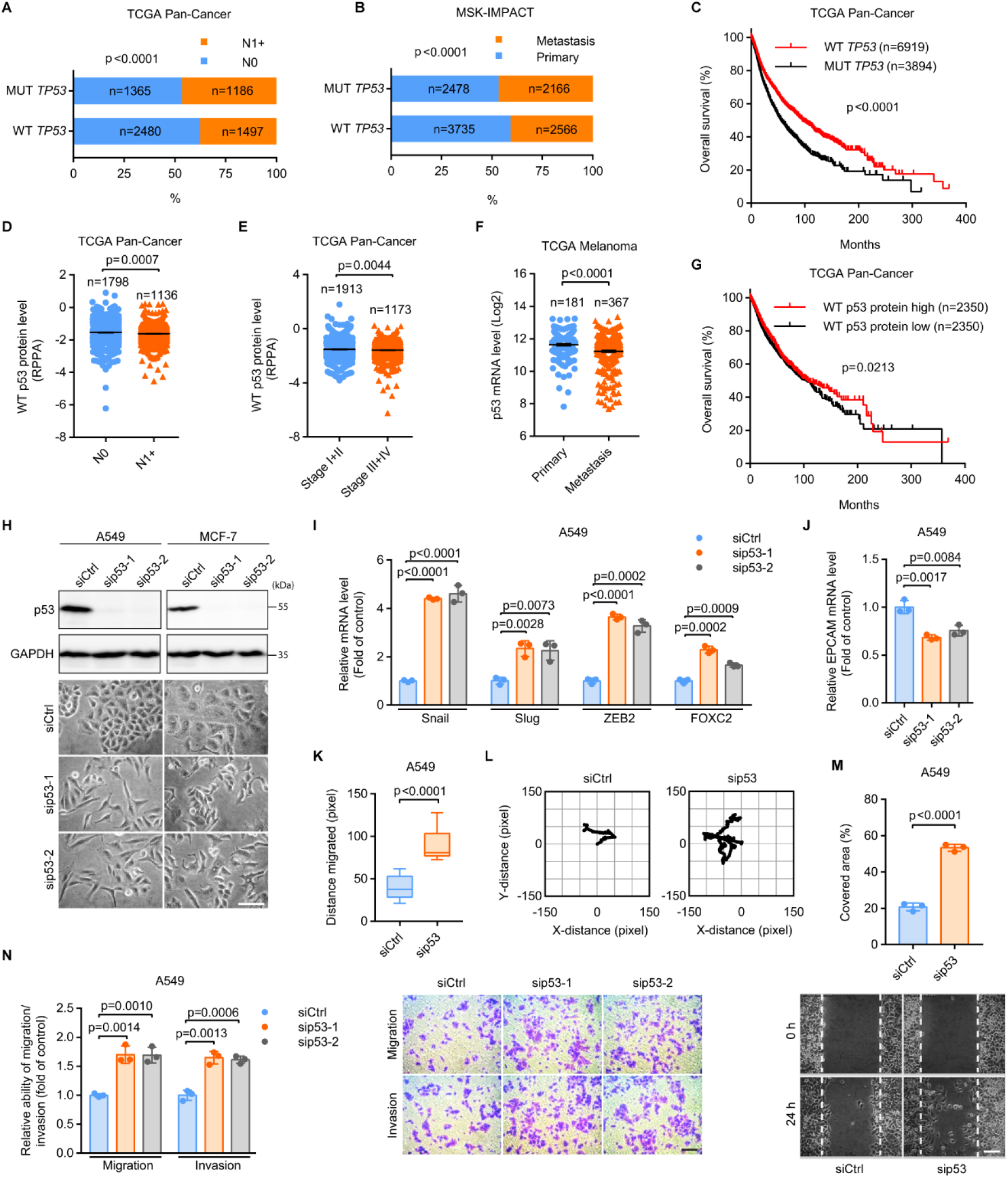
Downregulation of WT p53 expression is associated with aggressive tumor phenotypes and poor prognosis. **A**, **B** Contingency analysis of the associations between the presence of *TP53* mutations and the probabilities of metastases to (**A**) lymph nodes and (**B**) distant organs. N0, lymph node-negative; N1+, lymph-node-positive. Data were derived from (**A**) The Cancer Genome Atlas (TCGA) Pan-Cancer and (**B**) the Memorial Sloan-Kettering Integrated Mutation Profiling of Actionable Cancer Targets (MSK-IMPACT) cohorts. **C** Kaplan-Meier analysis of overall survival in cancer patients having WT and MUT *TP53*. **D**, **E** p53 protein expression of patients having WT *TP53* (**D**) with N0 and N1+ and (**E**) with stage I+II and III+IV tumors. RPPA, reverse-phase protein array. **F** p53 mRNA levels in primary and distant metastatic melanoma. **G** Kaplan-Meier analysis of overall survival in *TP53* WT cancer patients with low and high p53 protein levels. Data were extracted from TCGA (**C**-**G**). **H** Phase-contrast imaging of control (siCtrl) and p53-silenced (sip53-1 and sip53-2) A549 and MCF-7 cells. Scale bar: 100 µm. **I**, **J** qRT-PCR analysis of the mRNA levels of (**I**) EMT inducers Snail, Slug, ZEB2, and FOXC2 and (**J**) the epithelial cell adhesion molecule EPCAM in siCtrl, sip53-1, and sip53-2 A549 cells. **K**, **L** Migration distance (**K**) and representative trajectories (**L**) of siCtrl (n = 28) and sip53 (n = 14) A549 cells. **M** Quantification (top) and representative images (bottom) of the area in a wound-healing assay covered by A549 cells transfected with siCtrl or sip53. Scale bar: 100 µm. **N** Transwell assays for siCtrl, sip53-1, and sip53-2 A549 cells. Scale bar: 100 µm. Error bars represent mean ± SEM (**D**-**F**) or SD (**I**, **J**, **M**, **N**). Data were analyzed by Fisher’s exact test (**A**, **B**), log-rank test (**C**, **G**), or two-tailed unpaired Student’s t test (**D**-**F**, **I**-**K**, **M**, **N**)

Further details for the materials and methods used in this study can be found in Supplementary Materials and Methods.

## Results

### Downregulation of WT p53 expression is associated with aggressive tumor phenotypes and poor prognosis

Given that *TP53* is among the most frequently altered genes in metastatic cancers [34], we investigated the associations between the presence of *TP53* mutations and cancer metastases using The Cancer Genome Atlas (TCGA) Pan-Cancer (Fig. 1A) and the Memorial Sloan-Kettering Integrated Mutation Profiling of Actionable Cancer Targets (MSK-IMPACT) (Fig. 1B) cohorts. Results revealed that cancer patients harboring mutant (MUT) *TP53* had a higher risk of developing metastases to lymph nodes (Fig. 1A, lymph node-negative (N0) vs lymph node-positive (N1+)) and distant organs (Fig. 1B) as compared to those having WT *TP53*. In addition, overall survival was significantly longer in patients with WT *TP53* than in those with MUT *TP53* (Fig. 1C). As emerging evidence demonstrates the important role of WT p53 in suppressing cancer metastasis [27], we reasoned that the expression level of WT p53 might also be a contributing factor in determining disease aggressiveness and patient outcomes. To this end, we analyzed the protein levels of p53 in primary tumors from N0 and N1+ cancer patients harboring WT *TP53* using the reverse-phase protein array (RPPA) data derived from the TCGA Pan-Cancer dataset (Fig. 1D). Results indicated that N1+ tumors had decreased protein levels of WT p53 as compared to those in N0 tumors. Of tumors with WT *TP53*, levels of p53 protein expression were also reduced in advanced-stage (III+IV) tumors when compared to those in the earlier-stage (I+II) tumors (Fig. 1E). In line with these observations, p53 mRNA levels were significantly decreased in distant metastatic compared to those in primary melanoma (Fig. 1F). Reduced p53 protein levels were strongly correlated with impaired overall survival in cancer patients harboring WT *TP53* (Fig. 1G). These data suggest that impaired expression of WT p53 is implicated in exaggerated malignant phenotypes and poor prognosis of cancer patients.

### p53 silencing accelerates cancer cell migration and invasion

To validate the contribution of WT p53 in suppressing cancer dissemination, we silenced p53 in human non-small cell lung cancer (NSCLC) A549 and human breast cancer MCF-7 cells with small interference (si)RNAs. Both cell types express WT p53. Phase-contrast imaging indicated that p53 depletion induced significant morphological changes in both A549 and MCF-7 cells (Fig. 1H). p53-depleted cells exhibited a decrease in cell-cell adhesions, an elongated cell body, and a spindle-shaped morphology, which were much different from the high cell-cell adhesion and epithelial-like morphology in p53 WT controls. Because the phenotypic switch from epithelial-to spindle-like morphology is one of the hallmarks of epithelial-to-mesenchymal transition (EMT), allowing cells to become motile and invasive, we investigated the role of p53 in regulating the expression of EMT signaling molecules and cell migration and invasion. As expected, p53 depletion elevates the mRNA levels of EMT-promoting factors Snail (Snail1), Slug (Snail2), ZEB2 (zinc finger E-box binding homeobox 2), and FOXC2 (forkhead box protein C2) (Fig. 1I) but repressed the epithelial cell adhesion molecule EPCAM (Fig. 1J) in A549 cells, suggesting that loss of p53 triggers EMT, a key event that drives cancer metastasis. Moreover, single-cell tracking and wound-healing assays showed that p53 silencing stimulated A549 cell motility (Fig. 1K-M). Transwell assays further confirmed that p53-silenced cells were more migratory and invasive than p53 WT controls and invasion (Fig. 1N). Taken together, these results indicate that loss of WT p53 induced a more aggressive cancer cell phenotype and heightened cell motility and invasion.

### p53 silencing amplifies mitochondrial fission has diagnostic and clinical implications

Our results highlight an important role of WT p53 in restraining the metastatic dissemination of cancer cells. Building on previous findings that metastasizing cancer cells need to alter their mitochondrial morphology to facilitate their motility and invasiveness [6, 9, 35], we assessed the morphological dynamics of mitochondria upon WT p53 loss. Using live-cell fluorescence imaging, we observed cells harboring WT p53 have predominantly intermediate mitochondria (>78%). Upon p53 silencing, cells with fragmented mitochondria were dramatically enhanced while cells with elongated and intermediate mitochondria were drastically reduced. Mitochondria were fragmented in more than 80% of p53-depleted cells (less than 2% in p53 WT controls) (Fig. 2A, B). These results provide strong evidence that mitochondrial dynamics is modulated by p53.

**Fig. 2.**
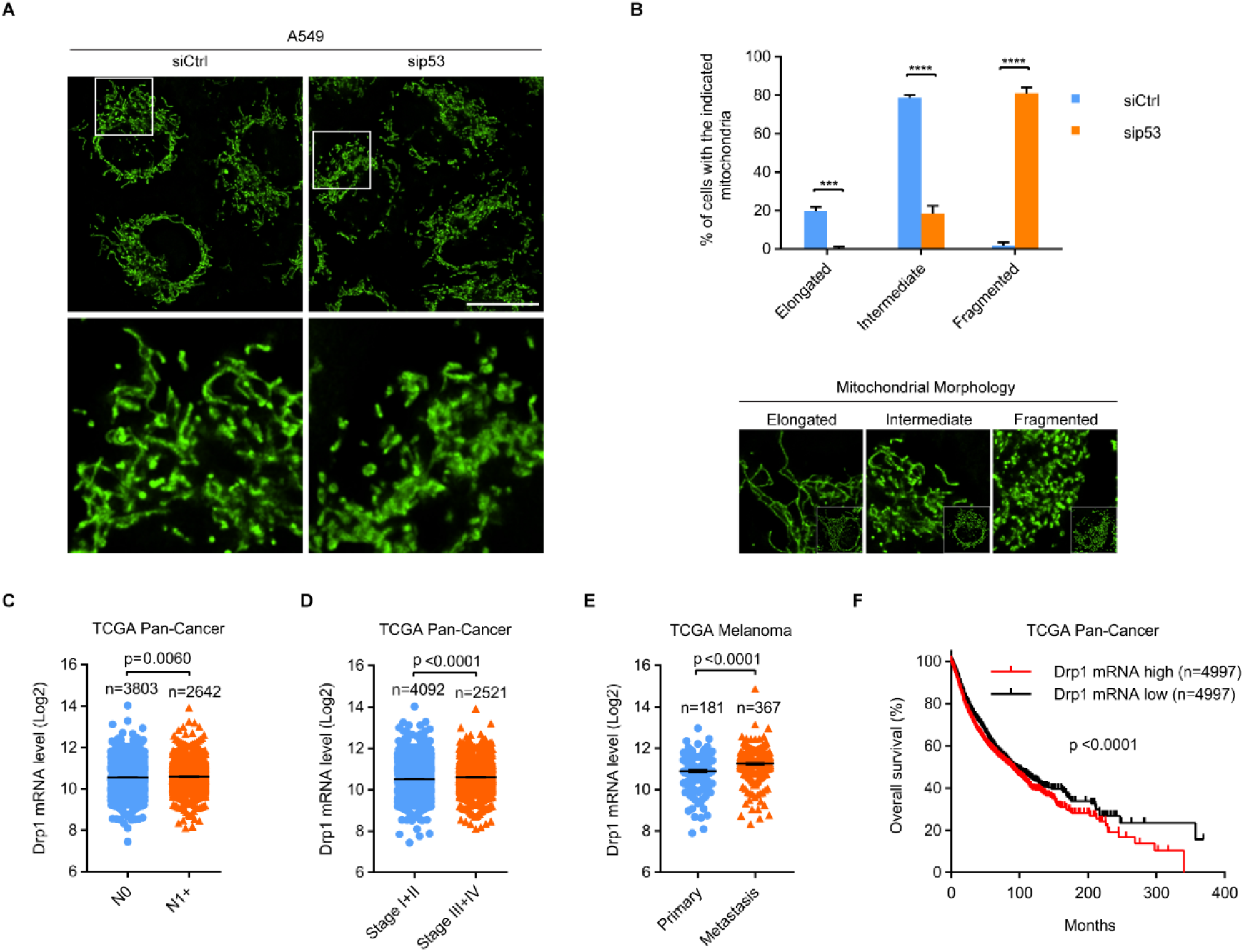
p53 silencing amplifies mitochondrial fission has diagnostic and clinical implications. **A**, **B** Representative images (**A**) and quantification (**B**) of mitochondrial morphology in siCtrl (n = 222) and sip53 (n = 230) A549 cells. Boxed regions in **A** are shown enlarged in the bottom panels. Scale bar: 20 µm. **B** Representative images for each mitochondrial morphology type are shown in the bottom panels. **C**, **D** Drp1 mRNA expression in (**C**) N0 and N1+ or (**D**) stage I+II and III+IV tumors. **E** Drp1 mRNA levels in primary and distant metastatic melanoma. **F** Kaplan-Meier analysis of overall survival in cancer patients with low and high Drp1 mRNA levels. Data were extracted from TCGA (**C**-**F**). Error bars represent mean ± SD (**B**) or SEM (**C**-**E**). Data were analyzed by two-tailed unpaired Student’s t test (**B**-**E**) or log-rank test (**F**). *\*\*\**, *p* < 0.001; *\*\*\*\**, *p* < 0.0001

As p53 silencing amplified mitochondrial fission, we investigated the clinical significance of Drp1, a major pro-fission protein. The mRNA levels of Drp1 were elevated in N1+ tumors compared to those in N0 tumors (Fig. 2C). Consistent with this finding, we also observed an increase in mRNA levels of Drp1 in advanced-stage (III+IV) tumors when compared to those in the earlier-stage (I+II) tumors (Fig. 2D). Especially, Drp1 mRNA levels were higher in distant metastatic than in primary melanoma (Fig. 2E). Furthermore, high Drp1 levels significantly reduced overall survival in cancer patients (Fig. 2F). These data corroborate a strong association of mitochondrial morphology with the degree of tumor malignancy and clinical outcomes in cancers.

### p53 elevation promotes mitochondrial elongation accompanied by attenuated invasive cell migration

Sodium arsenite (SA) is a genotoxic agent that induces DNA damages in human cell lines [36, 37] and elevates endogenous p53 expression (Fig. S1A). We examined the effects of SA-induced endogenous p53 upregulation on mitochondrial dynamics and the metastatic abilities of cancer cells. Although 10 µM SA was sufficient to induce p53 expression in A549 cells (Fig. S1A), treating cells with 10, 20, or 40 µM SA did not affect cell viability and proliferation (Fig. S1B, C). Furthermore, A549 cells treated with increased concentration (20 or 40 µM) of SA had higher accumulations of p53 protein. Thus, a 24-h treatment with a non-cytotoxic concentration of 20 µM SA was chosen to amplify the endogenous p53 expression in A549 cells. The expression of the endogenous p53 with and without SA induction can be effectively silenced with siRNA (Fig. 3A). Critically, these treatments did not change the survival of p53-depleted cells (Fig. S1D).

**Fig. 3.**
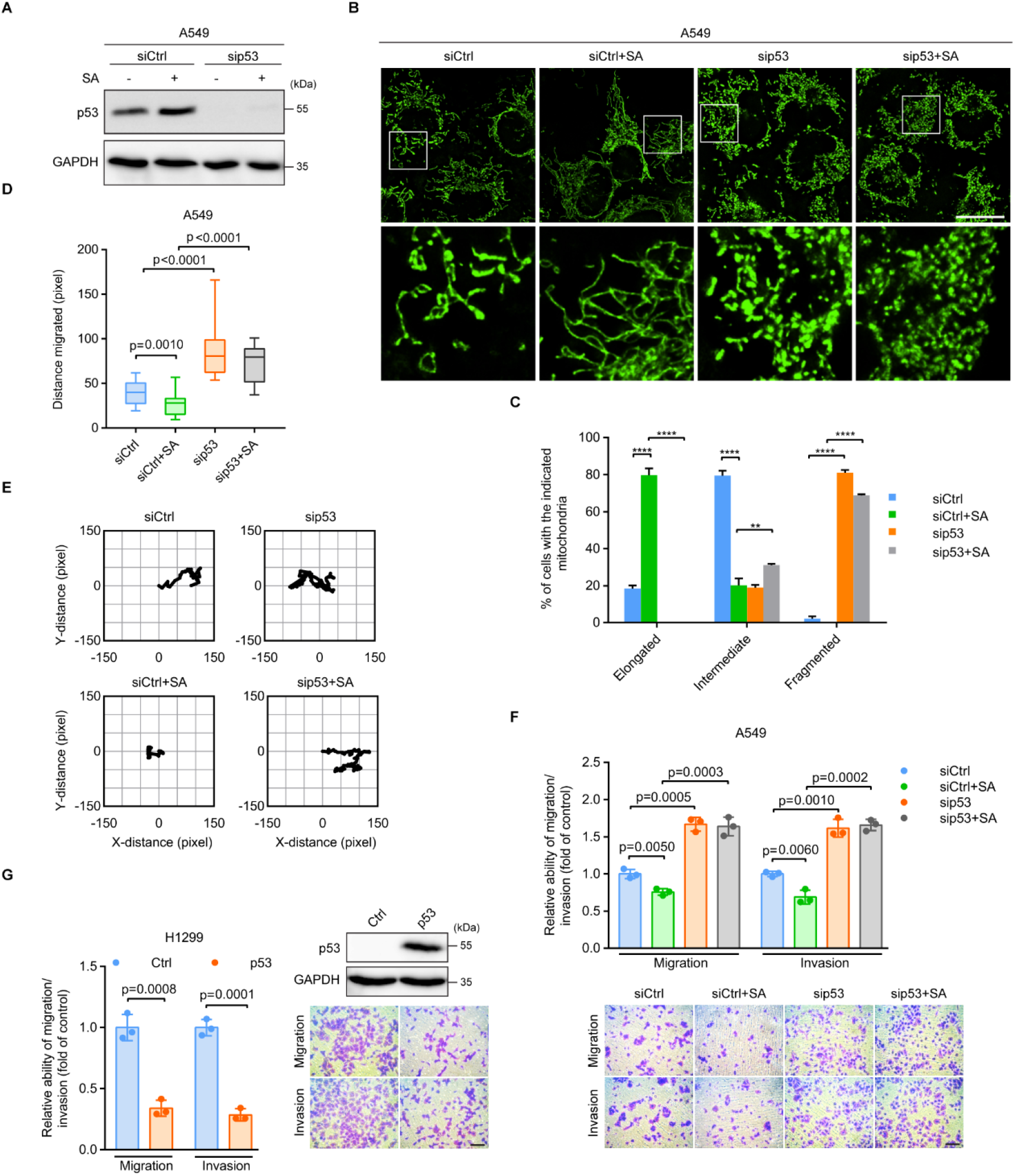
p53 elevation promotes mitochondrial elongation accompanied by attenuated invasive cell migration. **A** Immunoblot of p53 in siCtrl and sip53 A549 cells with and without 20 µM SA treatment for 24 h. GAPDH was used as a loading control. **B**, **C** Representative images (**B**) and quantification (**C**) of mitochondrial morphology in siCtrl (n = 223), siCtrl+SA (n = 211), sip53 (n = 205), and sip53+SA (n = 218) A549 cells. Boxed regions in **B** are shown enlarged in the bottom panels. Scale bar: 20 µm. **D**, **E** Migration distance (**D**) and representative trajectories (**E**) of siCtrl (n= 29), siCtrl+SA (n = 17), sip53 (n = 13), and sip53+SA (n = 15) A549 cells. **F**, **G** Transwell assays for (**F**) siCtrl, siCtrl+SA, sip53, and sip53+SA A549 cells or (**G**) control (Ctrl) and p53-overexpressing (p53) H1299 cells. Scale bar: 100 µm. Error bars represent mean ±SD. Data were analyzed by two-tailed unpaired Student’s t test. *\*\**, *p* < 0.01; *\*\*\*\**, *p* < 0.0001

We have shown that cells with p53 knockdown have decreased mitochondrial elongation and enhanced mitochondrial fragmentation (Fig. 2A, B). In contrast, SA-elevated endogenous p53 expression promoted a robust increase in mitochondrial branching and elongation (Fig. 3B, C). In SA-treated A549 cells, almost 80% of cells showed elongated mitochondria as compared to 20% in control cells. p53 depletion completely reversed the effects of SA on mitochondrial elongation, accompanied by enhanced mitochondrial fragmentation. Over 70% of SA-treated and p53-silenced cells exhibited fragmented mitochondria that were absent in SA-treated only p53 WT cells. Intriguingly, mitochondrial membrane potential (MMP) and levels of mitochondrial reactive oxygen species (ROS) were unaltered by increasing or decreasing the expression of p53 (Fig. S1E, F). These results underscore the central role of p53 in the control of mitochondrial dynamics without affecting mitochondrial integrity.

Notably, single-cell tracking results showed that SA-treated cells exhibited a 1.5-fold decrease in migratory capacity as compared to that of control cells (Fig. 3D, E). Knockdown of p53 fully abolished the inhibitory effects of SA on cell migration. p53 silencing accelerated the migratory ability of SA-treated cells to a level similar to that of untreated and p53-depleted cells. In line with this, transwell assays also showed that SA lowered cell migration and invasion but these effects were abolished by p53 silencing (Fig. 3F). These results further support the suppressive role of p53 in the metastatic properties of cancer cells.

To further substantiate the role of p53 in regulating cell migration and invasion, we transfected the pcDNA3 p53 WT plasmid into the p53-null NSCLC H1299 cells to express exogenous WT p53 (Fig. 3G). As monitored by wound-healing assays, expression of WT p53 caused an approximately 2.5-fold decrease in the migratory capacity of the cells as compared to that of the controls (Fig. S1G). Corroborating this finding, transwell assays also showed that expression of WT p53 significantly inhibited cell migration and invasion (Fig. 3G). In conclusion, these results, together with the observation that p53 depletion exaggerated mitochondrial fragmentation and invasive cell migration, pinpoint p53 as a potent regulator of mitochondrial dynamics and cell motility.

### p53 alleviates Drp1-mediated mitochondrial fission and thereby restrains cell migration and invasion

To delineate the underlying molecular mechanism responsible for enhanced mitochondrial fragmentation during p53 loss, we investigated changes in the expression and phosphorylation of mitochondrial fusion and fission factors in response to changes in p53 levels. Our qRT-PCR results showed that mRNA levels of mitochondrial fusion (Mfn1, Mfn2, and Opa1) and fission (Drp1, Fis1, Mff, and MIEF1) factors were unaltered upon p53 silencing in A549 cells (Fig. S2A). Additionally, both SA-induced p53 upregulation and siRNA-mediated p53 knockdown did not affect the protein levels of total Drp1 and Mfn2 (Fig. 4A).

**Fig. 4.**
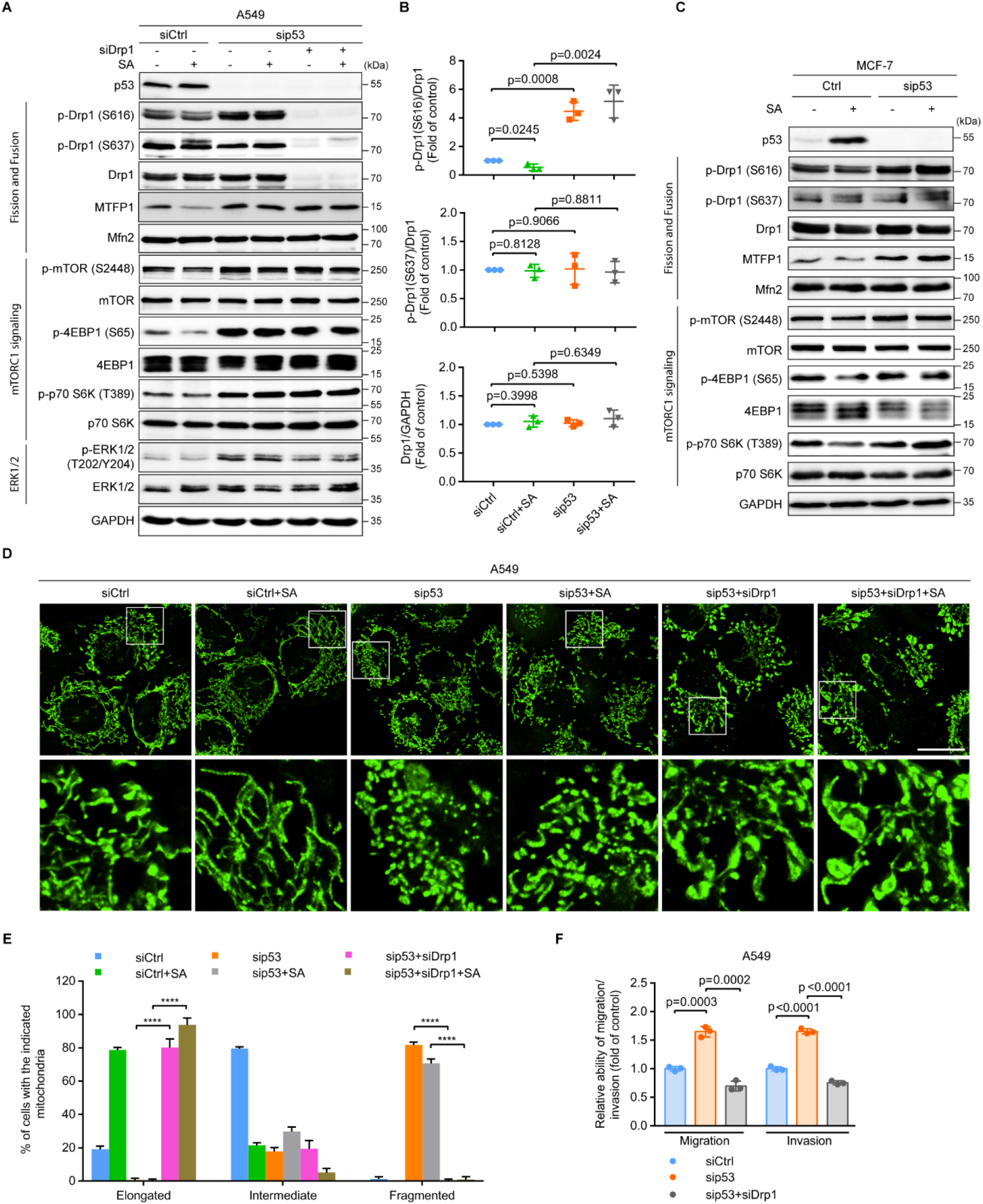
p53 alleviates Drp1-mediated mitochondrial fission and thereby restrains cell migration and invasion. **A** Immunoblot of the indicated proteins in siCtrl, sip53, and p53/Drp1 double-knockdown (sip53+siDrp1) A549 cells with and without 20 µM SA treatment for 24 h. GAPDH was used as a loading control. **B** Quantification of levels of Drp1, p-Drp1 (S637), and p-Drp1 (S616) in **A**. **C** Immunoblot of the indicated proteins in siCtrl and sip53 MCF-7 cells with and without 20 µM SA treatment for 24 h. GAPDH was used as a loading control. **D**, **E** Representative images (**D**) and quantification (**E**) of mitochondrial morphology in siCtrl (n = 226), siCtrl+SA (n = 232), sip53 (n = 205), sip53+SA (n = 214), sip53+siDrp1 (n = 230), and sip53+siDrp1+SA (n = 214) A549 cells. Boxed regions in **D** are shown enlarged in the bottom panels. Scale bar: 20 µm. **F** Transwell assays for siCtrl, sip53, and sip53+siDrp1 A549 cells. Error bars represent mean ± SD. Data were analyzed by two-tailed unpaired Student’s t test. *\*\*\*\**, *p* < 0.0001

Phosphorylation of serine 616 (S616) on Drp1 enables Drp1-directed mitochondrial fission, whereas phosphorylation of serine 637 (S637) on the same protein abolishes its GTPase activity and inhibits the fission of mitochondria [15]. While the levels of Drp1 S637 phosphorylation were unaffected by the expression levels of p53, SA-elevated p53 expression that caused mitochondrial elongation (Fig. 3B, C) triggered an ∼50% reduction in the pro-fission S616 phosphorylation of Drp1 (Fig. 4A, B). Conversely, in both untreated and SA-treated p53-silenced cells whose mitochondria were extensively fragmented (Fig. 3B, C), Drp1 S616 phosphorylation was increased by over 4-fold when compared to that of the p53 WT controls (Fig. 4A, B). These results suggest that p53 might control mitochondrial morphology by modulating the phosphorylation of S616 on Drp1.

Reportedly, mTORC1 phosphorylates 4EBPs to enable MTFP1 translation, thereby stimulating Drp1-mediated mitochondrial fission [17]. To illuminate how p53 controls Drp1 S616 phosphorylation, we examined the effects of p53 on the MTFP1 protein level and mTORC1 activity. In agreement with the Drp1 S616 phosphorylation and mitochondrial fission activity, the levels of MTFP1 and proteins relevant to mTORC1 signaling, including the phosphorylations of mTOR S2448, 4EBP1 S65, and S6K1 (p70 S6K) T389 were significantly decreased following SA-stimulated endogenous p53 expression in A549 cells. p53 depletion dramatically enhanced the MTFP1 protein level and mTORC1 activity in both untreated and SA-treated cells (Fig. 4A). Furthermore, elevated p53 expression by SA also hampered the T202/Y204 phosphorylation of ERK1/2 (Fig. 4A) whose activation was governed by Drp1-mediated mitochondrial fission [6]. p53 silencing, however, robustly augmented ERK1/2 phosphorylation in both untreated and SA-treated cells (Fig. 4A).

To verify the regulation of mitochondrial dynamics and mTORC1 signaling by p53 is not restricted to A549 cells, we examined mitochondrial fission and fusion factors and mTORC1 signaling in MCF-7 (Fig. 4C) and H1299 (Fig. S2B, C) cells. Similar to findings in A549 cells, both SA-induced endogenous p53 upregulation and siRNA-mediated knockdown of p53 had no significant effect on the levels of Drp1, Mfn2, and Drp1 S637 phosphorylation in MCF-7 cells. Conversely, there was a robust increase in the S616 phosphorylation of Drp1, correlated with an elevation of MTFP1 protein levels and mTOR, 4EBP1, and S6K1 phosphorylations in both untreated and SA-treated MCF-7 cells upon p53 silencing (Fig. 4C). In sharp contrast to p53 WT A549 and MCF-7 cells, SA treatment did not affect the levels of Drp1, Mfn2, and Drp1 S637 phosphorylation, but it induced an ∼2.5-fold increase in the levels of Drp1 S616 phosphorylation and a corresponding upregulation of MTFP1 protein levels and mTOR, 4EBP1, and S6K1 phosphorylations in p53-null H1299 cells (Fig. S2B, C). Importantly, overexpression of exogenous WT p53 in control and SA-treated H1299 cells diminished Drp1 S616 phosphorylation by more than 40% and 50%, respectively, whereas levels of total Drp1, Mfn2, and Drp1 S637 phosphorylation were unaffected (Fig. S2B, C). In line with this, phosphorylations of mTOR, 4EBP1, and S6K1 and the protein levels of MTFP1 were strongly decreased upon exogenous expression of WT p53 in both untreated and SA-treated H1299 cells (Fig. S2B). Altogether, these results support the notion that p53 drives mitochondrial elongation by inhibiting the phosphorylation of S616 on Drp1, accompanied by reducing MTFP1 protein levels and the mTORC1 activity.

To further investigate whether Drp1-mediated mitochondrial fission might contribute to the metastatic phenotype driven by p53 loss, we performed a double-knockdown of both p53 and Drp1 in A549 cells. In line with our conjecture that mTORC1-controlled MTFP1 protein translation is the upstream signaling that modulates Drp1 activity and mitochondrial fission [17], co-knockdown of Drp1 in p53-silenced cells did not affect the total amounts and phosphorylations of mTOR, 4EBP1, and S6K1 and the protein levels of MTFP1 as compared to those in cells with p53 knockdown alone in either the absence or presence of SA (Fig. 4A). In contrast, ERK1/2 phosphorylation was strongly reduced in cells with p53/Drp1 double-knockdown when compared to that in cells with p53 knockdown alone (Fig. 4A), suggesting that ERK1/2 might be the downstream signaling regulated by Drp1-driven mitochondrial fission. Significantly, Drp1 depletion not only rescued p53 deficiency-induced mitochondrial fragmentation but also exaggerated mitochondrial elongation. Over 80% and 90% of mitochondria were elongated in untreated and SA-treated p53/Drp1 double-knockdown cells, respectively (as compared to <1% in untreated and SA-treated p53-depleted cells) (Fig. 4D, E). Most notably, Drp1 depletion abolished accelerated cell migration in both untreated and SA-treated p53 knockdown cells, as illustrated by single-cell tracking (Fig. S2D, E) and wound-healing assays (Fig. S2F). Consistently, transwell assays showed a more than 2-fold decrease in the migratory and invasive abilities of p53/Drp1 double-knockdown cells, when compared to those with only p53-knockdown (Fig. 4F). Taken together, these results demonstrate that elevated mitochondrial fragmentation caused by increased Drp1 S616 phosphorylation is responsible for the aggressive cell migration and invasion seen upon p53 loss.

### p53 diminishes mTORC1-controlled MTFP1 protein levels to attenuate Drp1-driven mitochondrial fission and invasive cell migration

Our results indicate that p53 silencing triggered mitochondrial fragmentation accompanied by an increase in Drp1 S616 phosphorylation and activation of mTORC1 signaling, whereas Drp1 depletion had no effect on the levels of mTORC1-relevant factors (Fig. 4A). Possibly, mTORC1 functions upstream of Drp1 to drive Drp1 S616 phosphorylation and is inhibited by p53. To this end, we next examined the relationship between p53 and mTORC1. As expected, Pan-Cancer database analysis showed that p53 protein levels correlated inversely with 4EBP1 S65, 4EBP1 T37/T46, and mTOR S2448 phosphorylations (Fig. 5A), suggesting a negative effect of p53 on mTORC1 activity. Since p53 acts predominantly as a transcription factor, we examined whether the transcriptional regulatory function of p53 is involved in the p53-driven suppression of mTORC1 activity and Drp1 S616 phosphorylation. Interestingly, the p53-specific transcriptional inhibitor, pifithrin-α (PFT-α), successfully suppressed the induction of the p53 downstream target p21 under the condition of SA-induced p53 upregulation. Furthermore, PFT-α did not affect p53 protein levels, but strongly enhanced mTOR, 4EBP1, and Drp1 S616 phosphorylations and the MTFP1 protein levels in both control and SA-treated A549 cells (Fig. S3A). These results are in line with the observations using p53 gene knockdown, suggesting that the transcriptional activity of p53 is essential for p53-mediated inhibition of mTORC1 and Drp1 activities. Remarkably, p53 silencing attenuated the gene expression of negative regulators of mTOR signaling, including PTEN, AMPKβ1, Sestrin1/2, and TSC2 (Fig. S3B).

**Fig. 5.**
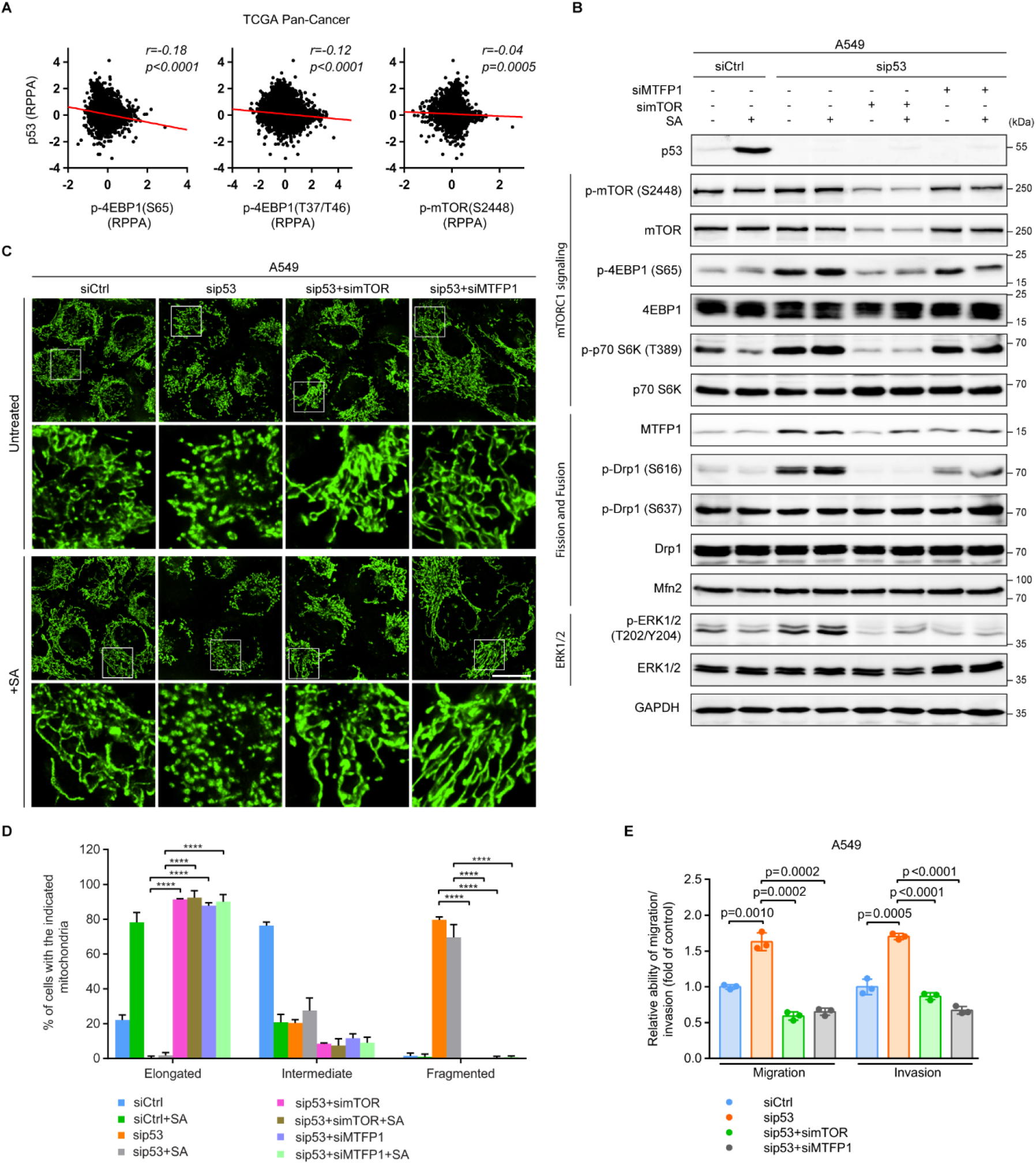
p53 diminishes mTORC1-controlled MTFP1 protein levels to attenuate Drp1-driven mitochondrial fission and invasive cell migration. **A** Correlations between the RPPA levels of p53 and the indicated proteins (n = 7 694 samples). Data were extracted from TCGA. **B** Immunoblot of the indicated proteins in siCtrl, sip53, p53/mTOR double-knockdown (sip53+simTOR), and p53/MTFP1 double-knockdown (sip53+siMTFP1) A549 cells with and without 20 µM SA treatment for 24 h. GAPDH was used as a loading control. **C**, **D** Representative images (**C**) and quantification (**D**) of mitochondrial morphology in siCtrl (n = 203), siCtrl+SA (n = 207), sip53 (n = 200), sip53+SA (n = 213), sip53+simTOR (n = 234), sip53+simTOR+SA (n = 209), sip53+siMTFP1 (n = 214), and sip53+siMTFP1+SA (n = 213) A549 cells. Boxed regions in **C** are shown enlarged in the bottom panels of each group. Scale bar: 20 µm. **E** Transwell assays for siCtrl, sip53, sip53+simTOR, and sip53+siMTFP1 A549 cells. Error bars represent mean ± SD. Data were analyzed by two-tailed unpaired Student’s t test. *\*\*\*\**, *p* < 0.0001

Increased or decreased p53 expression did not alter MTFP1 mRNA levels (Fig. S3C), suggesting that MTFP1 expression was controlled at the protein level. mTORC1 signaling reportedly mediates MTFP1 protein translation to govern Drp1 S616 phosphorylation and thereby mitochondrial fission [17]. To experimentally verify the interplay and chronology of mTORC1 signaling, MTFP1, and Drp1, we performed double knockdowns of p53 and mTOR or p53 and MTFP1 in A549 cells (Fig. 5B). Knockdown of mTOR reduced the phosphorylations of mTORC1 downstream effectors 4EBP1 and S6K1 and diminished MTFP1 protein levels in both untreated and SA-treated p53-silenced A549 cells. In contrast, MTFP1 knockdown did not affect the total amounts and phosphorylations of mTORC1-relevant factors. The mTOR or MTFP1 silencing successfully abolished increased Drp1 S616 phosphorylation in both untreated and SA-treated p53 knockdown cells, while both mTOR and MTFP1 knockdown did not affect total Drp1 and Mfn2 protein levels (Fig. 5B). In line with the reduction of ERK1/2 activity seen upon Drp1 silencing (Fig. 4A), ERK1/2 phosphorylation was dramatically diminished in both untreated and SA-treated p53/mTOR or p53/MTFP1 double-knockdown cells when compared with similar cells with only p53 silencing (Fig. 5B). These results provide compelling evidence that mTORC1 mediating MTFP1 protein expression is required for the increased Drp1 S616 phosphorylation during p53 loss.

We further unraveled the contribution of the mTORC1/MTFP1 axis in modulating mitochondrial dynamics. Knockdown of either mTOR or MTFP1 completely rescued fragmented mitochondria in both untreated and SA-treated p53-depleted cells (Fig. 5C, D). Mitochondria were elongated in ∼90% of p53/mTOR or p53/MTFP1 double-knockdown cells untreated or treated with SA (Fig. 5D). Consistent with the finding that mitochondrial dynamics impacts cell migration and invasion, exaggerated mitochondrial elongation triggered by mTOR or MTFP1 knockdown fully suppressed the migratory capacity of both untreated and SA-treated p53-depleted cells (Fig. S3D, E). Transwell assays confirmed the dramatic reduction of both migratory and invasive abilities in p53/mTOR or p53/MTFP1 double-knockdown cells as compared to those in cells with p53 silenced alone (Fig. 5E). In conclusion, these results demonstrate that the control of MTFP1 protein levels by mTORC1 signaling is critical for p53 deficiency-induced mitochondrial fragmentation and accelerated cell migration and invasion.

### Activation of mTORC1/MTFP1/Drp1/ERK1/2 signaling axis is required for the EMT switch, MMP9 elevation, and cancer dissemination upon WT p53 loss

Building on our findings that p53 silencing elevated ERK1/2 phosphorylation but could be counteracted by co-knockdown of Drp1, mTOR, or MTFP1 (Fig. 4A and 5B), we theorized that mitochondrial fission governed by mTORC1/MTFP1/Drp1 axis might activate ERK1/2 signaling to direct cell migration and invasion upon p53 loss. Indeed, we observed that ERK1/2 phosphorylation displayed inverse and direct correlations with p53 and the phosphorylations of mTORC1-relevant factors (mTOR and 4EBP1), respectively (Fig. 6A). Inhibition of ERK1/2 with PD98059 had no significant effect on MTFP1 protein levels and the phosphorylations of Drp1 and 4EBP1 in p53-silenced cells with and without SA-treatment (Fig. 6B). In contrast, similar to observations seen upon depletion of Drp1, mTOR, or MTFP1, inhibition of ERK1/2 rescued the aggressive cell phenotype caused by p53 silencing (Fig. S4A), diminished the expression of the EMT-promoting transcription factor Snail (Fig. 6C), and restored EPCAM mRNA levels (Fig. 6D). ERK1/2 inhibition also hampered p53 deficiency-accelerated cell motility and invasion (Fig. 6E and S4B, C). Thus, ERK1/2 signaling functions downstream of mitochondria whose morphology is modulated by the mTORC1/MTFP1/Drp1 axis to favor cell dissemination upon p53 loss.

**Fig. 6.**
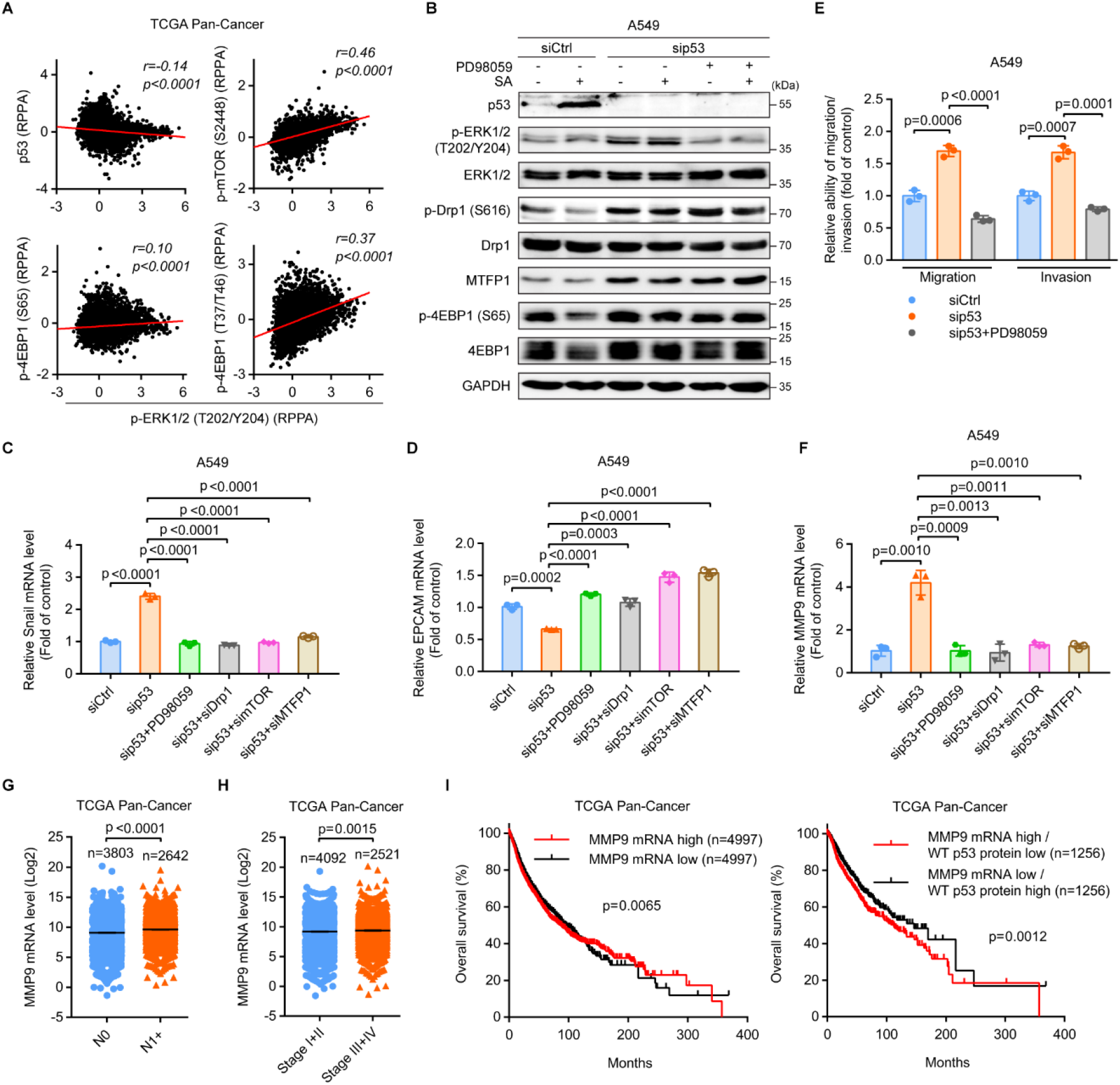
Activation of mTORC1/MTFP1/Drp1/ERK1/2 signaling axis is required for the EMT switch, MMP9 elevation, and cancer dissemination upon WT p53 loss. **A** Correlations between the RPPA levels of p-ERK1/2 (T202/Y204) and the indicated proteins (n = 7 694 samples). **B** Immunoblot of the indicated proteins in siCtrl, sip53, and PD98059-treated sip53 (sip53+PD98059) A549 cells with and without 20 µM SA treatment for 24 h. GAPDH was used as a loading control. **C**, **D** qRT-PCR analysis of the mRNA levels of (**C**) Snail and (**D**) EPCAM in siCtrl, sip53, sip53+PD98059, sip53+siDrp1, sip53+simTOR, and sip53+siMTFP1 A549 cells. **E** Transwell assays for siCtrl, sip53, and sip53+ PD98059 A549 cells. **F** qRT-PCR analysis of MMP9 mRNA expression in siCtrl, sip53, sip53+PD98059, sip53+siDrp1, sip53+simTOR, and sip53+siMTFP1 A549 cells. **G**, **H** MMP9 mRNA expression in (**G**) N0 and N1+ or (**H**) stage I+II and III+IV tumors. **I** Kaplan-Meier analysis of the overall survival in cancer patients with low and high MMP9 mRNA expression levels (left). Patients were further divided into MMP9 mRNA high / WT p53 protein low and MMP9 mRNA low / WT p53 protein high groups (right). Data were extracted from TCGA (**A**, **G**-**I**). Error bars represent mean ± SD (**C**-**F**) or SEM (**G**, **H**). Data were analyzed by two-tailed unpaired Student’s t test (**C**-**H**) or log-rank test (**I**)

Several studies have highlighted that ERK1/2 activation stimulates the expression of MMP9, which contributes to the proteolytic degradation of the extracellular matrix [38–41]. Strikingly, we observed elevated levels of MMP9 gene expression in p53-silenced A549 and MCF-7 cells when compared to those in p53 WT cells (Fig. 6F and S4D). Inhibition of ERK1/2 activation completely abolished increased MMP9 expression in p53-depleted cells. In line with this, co-knockdown of Drp1, mTOR, or MTFP1 in p53-silenced cells restored MMP9 expression to levels similar to the controls. Moreover, the levels of MMP9 mRNA were lower in cancer patients harboring WT *TP53* than in those with MUT *TP53* (Fig. S4E). The protein levels of p53 also exerted a negative correlation with the mRNA levels of MMP9 in cancers having WT *TP53* (Fig. S4F).

Consistent with previous literature showing that increased MMP9 expression promotes cancer cell migration, invasion, and metastasis [42], MMP9 mRNA levels were enhanced in N1+ (Fig. 6G) and advanced-stage (III+IV) (Fig. 6H) tumors when compared to those in N0 and the earlier-stage (I+II) tumors, respectively. MMP9 mRNA levels were also higher in metastatic than in primary melanoma (Fig. S4G). Higher MMP9 levels were associated with worse overall survival in cancer patients (Fig. 6I, left). The median overall survival was 86.14 and 93.20 months in patients expressing high and low MMP9 mRNA levels, respectively. Of cancer patients harboring WT *TP53*, those with high MMP9 mRNA and low p53 protein expression showed a significantly increased risk of death when compared to those with low MMP9 mRNA and high p53 protein expression (the median overall survival was 112.1 and 146.1 months, respectively) (Fig. 6I, right). Collectively, these results suggest that in p53-silenced cells, the mTORC1/MTFP1/Drp1/ERK1/2 axis might induce MMP9 expression to accelerate the metastatic dissemination of cancer cells and confirm the clinical implications for MMP9 in cancers.

## Discussion

This study unravels a molecular explanation of how tumor suppressor p53 modulates mitochondrial dynamics to restrict cancer cell dissemination. For the first time, we link p53 to mitochondria-dependent regulation of malignant properties of cancers, including cell motility and metastasis. p53 is canonically known to suppress cancer development by regulating multiple cell fate-determining genes involved in cell cycle arrest, DNA damage repair, senescence, and apoptosis [43]. p53 also constrains the metastatic abilities of cancer cells by transcriptionally controlling components of the metastatic cascade [27]. We present evidence here that p53 suppresses cancer dissemination via mitochondrial dynamics modulation and provide an alternative mechanism that coordinates aggressive phenotypes in cancers harboring compromised p53. The strong positive correlation between p53 expression levels and overall survival in cancer patients may be at least in part resulted from its anti-metastatic effects.

mTORC1 is commonly hyper-activated in p53-compromised cancers [44] and contributes to cancer cell migration [19, 20]. However, the underlying molecular signaling pathways remain poorly understood. We show here that mTORC1 accelerates cancer dissemination by directing MTFP1 protein expression. MTFP1 then facilitates Drp1-mediated mitochondrial fission upon p53 loss. These findings provide an important insight into a hitherto unknown mechanism that links mTORC1 to cancer metastasis. Recently, MTFP1 has also been implicated in promoting MMP9 expression and cancer metastasis although the underlying mechanism is still unknown [45]. Our results further corroborate the pro-migratory functions of MTFP1 and provide a molecular explanation for the observations. Accordingly, MTFP1 facilitates Drp1-mediated mitochondrial fission to enable ERK1/2 activation, thereby triggering the EMT-associated morphologic switch, MMP9 expression, and cancer cell dissemination. This underscores components of the mTORC1/MTFP1/Drp1/ERK1/2 signaling as potential and effective therapeutic targets for treating malignant and metastatic p53-compromised tumors [46, 47].

It is interesting to note that enhanced mitochondrial fragmentation is not always due to the accumulation of damaged mitochondria. Instead, mitochondrial biogenesis, by which new functional mitochondria are generated, also requires the initiation of Drp1-driven mitochondrial fission [48–50]. Fissions derived from mitochondrial dysfunction are associated with increased mitochondrial ROS and diminished MMP [49, 51, 52], whereas the mitochondrial physiology during fissions in the biogenesis of new mitochondria remains unchanged [49]. Intriguingly, our data show that depletion of p53 exaggerates mitochondrial fragmentation, but it does not affect MMP and mitochondrial ROS levels. In addition, it has been reported that mitochondrial biogenesis is regulated by the mTORC1/4EBP pathway which stimulates the translation of mRNAs encoding mitochondria-related proteins [21]. Here, we show that p53 depletion activates mTORC1/4EBP1 signaling that regulates MTFP1 protein expression to govern Drp1-mediated mitochondrial fission. Thus, we speculate that increased mitochondrial fission upon p53 loss is associated with stimulation of mitochondrial biogenesis, but not accumulation of damaged mitochondria. This would explain how the mitochondrial physiology remains constant in the context of p53 deficiency-induced mitochondrial fragmentation.

Accumulating evidence illustrates the critical roles of intracellular calcium (Ca^2+^) signaling in the regulation of key steps of the metastatic cascade, including EMT, focal adhesion turnover, lamellipodia formation, and the degradation of the extracellular matrix [53–55]. Notably, mitochondrial fission reduces the potential of endoplasmic reticulum (ER)-mitochondrial contacts and thereby attenuates the capacity of mitochondria to sequester Ca^2+^ released from the ER, leading to an increase in cytosolic Ca^2+^ levels [56, 57]. Moreover, mitochondrial fission resulting in elevated Ca^2+^ levels in the cytoplasm activates multiple Ca^2+^-dependent pathways regulating cellular behaviors, including cell migration and invasion [6, 58, 59]. Consistently, we find that exaggerated mitochondrial fission upon p53 loss triggers increased phosphorylation of ERK1/2, which is a downstream target of Ca^2+^/calmodulin-dependent protein kinase II (CaMKII), a major decoder of the intracellular Ca^2+^ oscillations [6, 60]. Intriguingly, pharmacological inhibition of ERK1/2 activity highlights an indispensable role of ERK1/2 signaling in controlling EMT, MMP9 expression, and the migratory and invasive abilities of cancer cells upon p53 loss. Our observations are supported by studies indicating that ERK1/2 stimulates MMP9 expression via regulating the activity of the transcription factors NF-ĸB (nuclear factor kappa-light-chain-enhancer of activated B cells) and AP-1 (activator protein-1) [40, 61, 62]. Furthermore, ERK1/2 signaling is implicated in controlling numerous other components of the cell motility machinery [63, 64]. For example, ERK1/2 signaling promotes cancer cell migration, invasion, and EMT by mediating the expression or the transcriptional activity of EMT-inducing transcription factors Twist1 [65, 66], Snail [67, 68], and Slug [69]. Thus, specific ERK1/2 inhibition may be beneficial to slow cancer metastasis in patients harboring compromised p53.

In summary, we have illustrated how p53 can modulate mitochondrial dynamics via controlling the mTORC1/MTFP1/Drp1 axis to restrict cancer cell dissemination (Fig. 7). Accordingly, cells with WT p53 exhibit basal mTORC1 activity, basal MTFP1 protein levels, and a coordinated balance between mitochondrial fission and fusion events. This maintains the mitochondrial morphology in a predominantly intermediate state and thereby prevents ERK1/2-governed cell migration and invasion. Cells with p53 silenced have increased activity of mTORC1 and the corresponding augmented MTFP1 protein levels. MTFP1 favors the pro-fission S616 phosphorylation of Drp1, which subsequently activates ERK1/2 signaling to enable EMT-associated morphological changes, MMP9 expression, and invasive cell migration. The molecular mechanism uncovered in this study is likely a general phenomenon. Indeed, we could observe the downregulation of WT p53 and the elevations of the pro-fission factor Drp1 and the metastatic driver MMP9 in aggressive and malignant tumors across cancers in The Cancer Genome Atlas. Additionally, Pan-Cancer analysis also showed that reduced WT p53 expression and enhanced expression of Drp1 or MMP9 are strongly correlated with poor prognosis. Our results offer a new molecular explanation for the aggressive malignant phenotypes of p53-compromised cancers and suggest that targeting mitochondrial fission and its downstream ERK1/2 signaling pathway may diminish the spread of p53-compromised cancer cells.

**Fig. 7.**
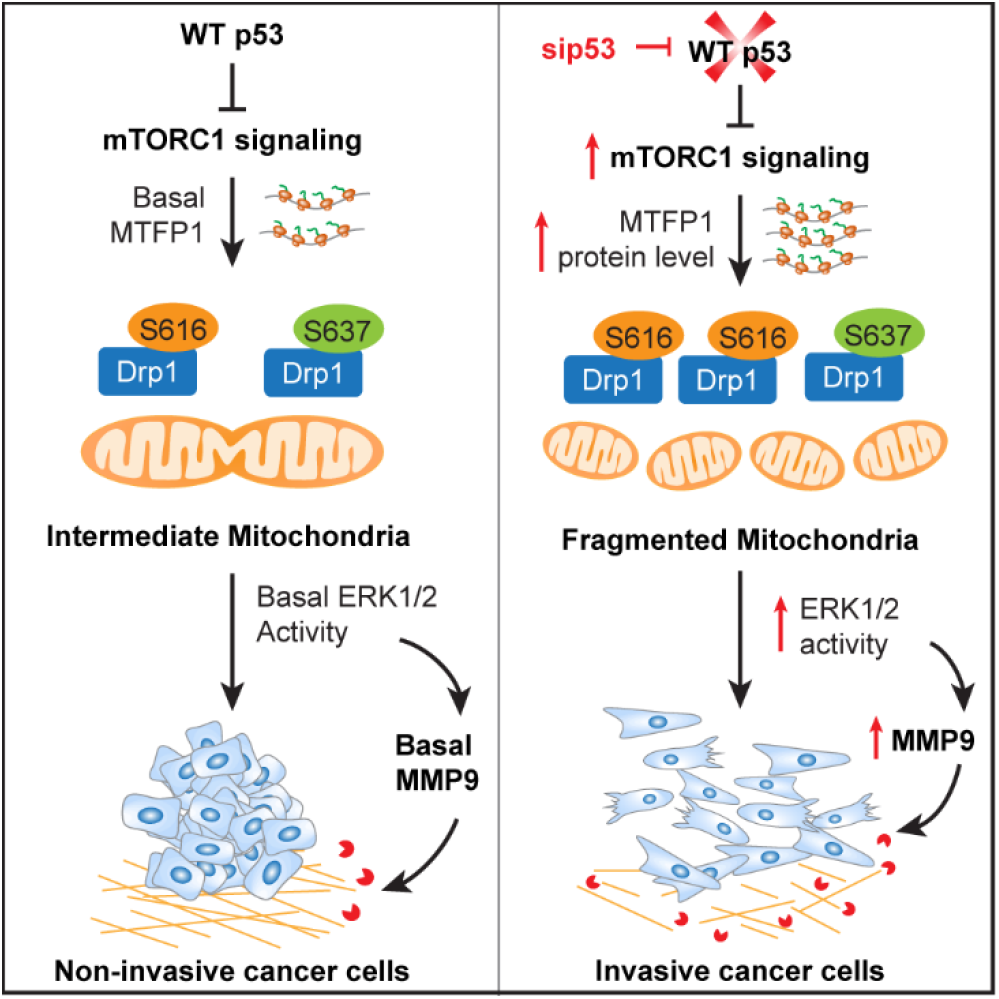
Schematic model of how p53 modulates mitochondrial dynamics to constrain EMT, MMP9 expression, and invasive cell migration. WT p53 suppresses mTORC1-directed MTFP1 protein expression and the aberrant phosphorylation of Drp1 at the pro-fission site S616, maintaining the predominantly intermediate state of mitochondria, and thereby constraining ERK1/2-mediated cell migration and invasion. Loss of WT p53 elevates mTORC1 activity, MTFP1 protein levels, and the phosphorylation of S616 on Drp1, shifting mitochondrial dynamics toward fission to promote ERK1/2 activation and resulting in EMT-like changes in cell morphology, increased MMP9 expression, and cell dissemination

## Statements & Declarations

### Acknowledgments

We thank Dr. Ming F. Tam (Department of Biological Sciences, Carnegie Mellon University, Pittsburgh, PA, USA) for the critical reading of the manuscript.

### Author Contributions

LYL and TTTP designed research; LYL, TTTP, and YCL performed research; TTTP and YCL analyzed data; YCL and YTC contributed analytic tools; CWW provided technical assistance; and LYL and TTTP wrote the paper.

### Data Availability

All data generated during this study are available from the corresponding author on reasonable request.

