## Supplementary Information for "Tumor Suppressor p53 Restrains Cancer Cell Dissemination by Modulating Mitochondrial Dynamics"

**This PDF file includes:**

Supplementary Materials and Methods

Figures S1 to S4

Tables S1 to S2

References for Supplementary Information

### Supplementary Materials and Methods

#### Cell viability assay

Cells were seeded in 96-well cell culture plates (TPP Techno Plastic Products AG, Trasadingen, Switzerland) at a density of $4×10^3$ cells per wells. Cell viability was assessed using MTT assay (Thermo Fisher Scientific) according to the manufacturer's instructions. Cells were added with 0.4 mg/ml of MTT reagent and incubated for 4 h at 37 °C before the absorbance at 550 nm was measured with a microtiter plate reader (Bio-Rad).

#### Colony formation assay

$1×10^3$ cells were seeded in 6-well plates (Corning) and incubated with complete medium at 37 °C in 5% $CO_2$ for 24 h to facilitate their attachment. Subsequently, cells were treated with various concentrations of sodium arsenite for an additional 24 h period before the medium was removed and replaced with the fresh medium containing 10% FBS. After 14 days of incubation, cells were fixed in 4% PFA (Electron Microscopy Sciences) for 20 min and stained with 0.05% crystal violet (Sigma-Aldrich) for 2 h. Colonies were imaged and measured by the ImageJ software (NIH, Bethesda, MD, USA).

#### Flow cytometry

All experiments were performed using 6-well plates (Corning). A total of 10,000 events, excluding debris, were recorded for each sample. All flow cytometric data were obtained using BD Accuri C6 flow cytometer (BD Biosciences, San Jose, CA, USA) and analyzed by the FlowJo 7.6.1 software (FlowJo LLC, Ashland, OR, USA).

For measurement of mitochondrial membrane potential (MMP) and mitochondrial reactive oxygen species (ROS), cells were treated with trypsin and prepared as single-cell suspensions. Cells were then stained either with 2 μM of the MMP probe JC-1 (Invitrogen) for 20 min or with 5 μM of the mitochondrial superoxide indicator MitoSOX Red (Invitrogen) for 30 min following the manufacturer's protocols. Cells were washed twice in ice-cold PBS before analysis with flow cytometry.

#### ***TP53*** **gene statuses and clinical correlations**

Data of *TP53* gene statuses (WT and MUT *TP53*) used to analyze the associations between the presence of TP53 mutations and the probabilities of metastases to lymph nodes (Fig. 1A) and distant organs (Fig. 1B) were derived from the TCGA Pan-Cancer and the Memorial Sloan-Kettering Integrated Mutation Profiling of Actionable Cancer Targets (MSK-IMPACT) [1] cohorts respectively, downloaded from the cBioPortal for Cancer Genomics (https://www.cbioportal.org/) [2, 3]. In each dataset, tumors were stratified into one of two categories: tumors with no *TP53* alteration (WT *TP53*), or tumors with one or more *TP53* mutations (MUT *TP53*). The complete TCGA Pan-Cancer dataset includes a total of 10 967 tumors from 10 953 patients across 32 different cancer types. *TP53* mutation data are available from 10 960 tumors. Among these tumor samples, 6 528 tumors have data of lymph node metastatic status (Fig. 1A) and 10 813 tumors have data of overall survival (Fig. 1C). The complete MSK-IMPACT cohort includes 10 945 tumors from 10 336 patients having both data of *TP53* mutations and tumor sites (Fig. 1B). The significance of the associations between the presence of TP53 mutations and the probabilities of metastases to lymph nodes (Fig. 1A) and distant organs (Fig. 1B) was determined by Fisher's exact test. The significance of the difference in the Kaplan-Meier plot of the overall survival in patients with WT and MUT *TP53* (Fig. 1C) was determined by log-rank (Mantel-Cox) test.

#### **Differential protein expression analysis**

Comparative analyzes of p53 protein expression levels (Fig. 1D, E) were performed using the reverse-phase protein array (RPPA) data acquired from the TCGA Pan-Cancer dataset in the cBioPortal for Cancer Genomics (https://www.cbioportal.org/) [2, 3]. A total of 4 741 tumors harboring WT *TP53* and having RPPA data were retrieved. Of which, 2 934 tumors have data of lymph node metastatic status (Fig. 1D) and 3 086 tumors have data of disease stages (Fig. 1E). RPPA values for p53 expression levels were stratified according to the lymph node metastatic status and the stage of the corresponding patient tumor. Mean protein expression levels of p53 were determined in lymph node-negative (N0) and -positive (N1+) (Fig. 1D) or earlier-stage (stage I+II) and advanced-stage (stage III+IV) (Fig. 1E) groups and significant downregulation of p53 protein expression in N1+ and advanced-stage tumors were statistically analyzed by two-tailed Student's t test (unpaired).

**Differential gene expression analysis**

Comparative analysis of gene expression patterns of p53 (Fig. 1F), Drp1 (Fig. 2E), and MMP9 (Fig. S4G) between primary and metastatic melanoma was performed using the RNA-Seq by Expectation-Maximization (RSEM) data extracted from TCGA in The UCSC Xena Browser (http://xena.ucsc.edu/) [4]. Pan-Cancer analyzes of the differential gene expression of Drp1 (Fig. 2C, D) and MMP9 (Fig. 6G, H) among tumor samples obtained from patients with lymph node-negative (N0) and -positive (N1+) prognostics (Fig. 2C and 6G) or at different stages (stage I+II and III+IV) (Fig. 2D and 6H); and that of MMP9 (Fig. S4E) among tumor samples with WT and MUT *TP53* were performed using RSEM values acquired from the TCGA Pan-Cancer dataset in the cBioPortal for Cancer Genomics (https://www.cbioportal.org/) [2, 3]. RSEM data are available for 10 071 tumors. Of these tumors, 6 445 tumors have data of lymph node metastatic status (Fig. 2C and 6G), 6 613 tumors have data of cancer stages (Fig. 2D and 6H), and 10 070 tumors have data of *TP53* mutation status (Fig. S4E).

All RSEM values were log2 transformed. Similar to methods used for protein expression data, mean expression levels were determined for mRNAs in individual groups and significant up- or down-regulation of mRNA expression was statistically analyzed by two-tailed Student's t test (unpaired).

**Correlation analysis**

The correlation between protein expression levels of p53 versus mRNA expression levels of MMP9 (Fig. S4F) was determined using RPPA and RSEM values extracted from the TCGA Pan-Cancer dataset in the cBioPortal for Cancer Genomics (https://www.cbioportal.org/) [2, 3]. All RSEM values were log2 transformed. Of the 10 071 tumors with available RSEM data, 4 538 tumors harbor WT *TP53* and have RPPA data.

For the correlations between p53 protein expression levels versus 4EBP1 S65, 4EBP1 T37/T46, and mTOR S2448 phosphorylation levels (Fig. 5A); or ERK1/2 T202/Y204 phosphorylation levels versus p53 protein levels and phosphorylation levels of 4EBP1 S65, 4EBP1 T37/T46, and mTOR S2448 (Fig. 6A), we used level 4 normalized RPPA data of the TCGA Pan-Cancer cohort downloaded from The Cancer Proteome Atlas (https://tcpaportal.org/). The Pearson correlation coefficient (r) was used to establish the correlations between expression levels of proteins versus mRNAs, proteins versus proteins, and mRNAs versus mRNAs and determine the *p-value*.

**Survival analysis**

For correlation analysis of p53 protein expression levels and the overall survival of cancer patients harboring WT *TP53* (Fig. 1G), publicly available RPPA data and patients' overall survival status from the TCGA Pan-Cancer dataset were downloaded from the cBioPortal for Cancer Genomics (https://www.cbioportal.org/) [2, 3]. Of tumors with WT *TP53*, 4 700 tumors have both RPPA and overall survival data. Patients were split into high and low p53 expression groups based on the median values of p53 protein expression. The significance of the difference in the overall survival of patients with high and low p53 protein expression in Kaplan-Meier plots was determined by log-rank (Mantel-Cox) test.

For correlation analysis of mRNA expression levels of Drp1 (Fig. 2F) and MMP9 (Fig. 6I) versus overall survival of cancer patients, RSEM values and patients' overall survival data from the TCGA Pan-Cancer dataset were downloaded from the cBioPortal for Cancer Genomics (https://www.cbioportal.org/) [2, 3]. A total of 9 994 tumors having both RSEM and overall survival data were retrieved. Similar to methods used for the analysis of correlation of p53 protein expression patterns and overall survival, patients were split into high and low expression groups based on the median values of Drp1 and MMP9 mRNA expression. The significance of the difference in the overall survival of patients with high and low expressions of individual mRNAs in Kaplan-Meier plots was determined by log-rank (Mantel-Cox) test.

### Supplementary Figures

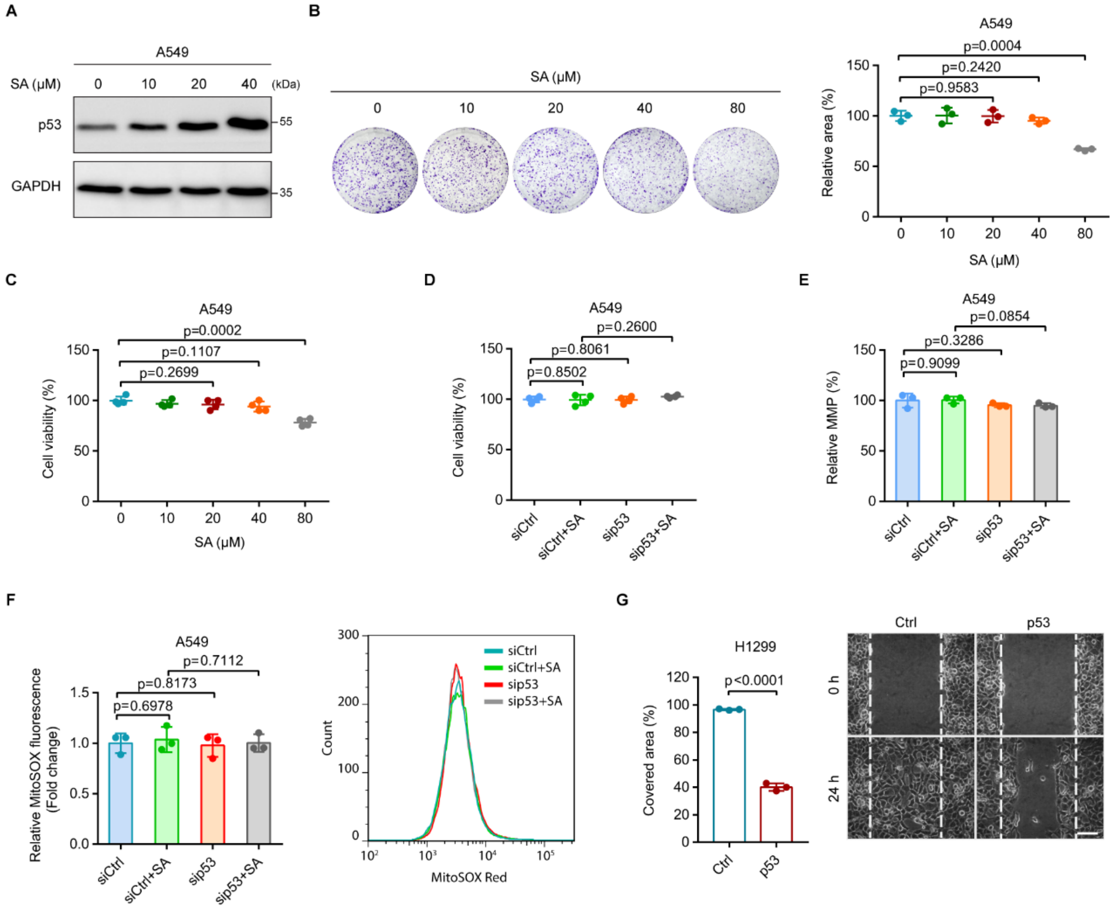


**Fig. S1** Effects of p53 on cell viability, mitochondrial function, and cell migration. **A** Immunoblot of p53 in A549 cells treated with SA at the indicated concentrations for 24 h. GAPDH was used as a loading control. **B**, **C** The viability of A549 cells treated with SA at the indicated concentrations for 24 h was measured by (**B**) colony formation assay or (**C**) MTT assay. **D** Cell viability was measured by MTT assay in siCtrl- and sip53-transfected A549 cells with and without 20 μM SA treatment for 24 h. **E**, **F** Flow cytometry analysis of (**E**) MMP or (**F**) mitochondrial ROS levels in siCtrl, siCtrl+SA, sip53, and sip53+SA A549 cells. **G** Quantification (left) and representative images (right) of the area in a wound-healing assay covered by Ctrl and p53-expressing H1299 cells. Scale bar: 100 μm. Error bars represent mean ± SD. Data were analyzed by two-tailed unpaired Student's t test.

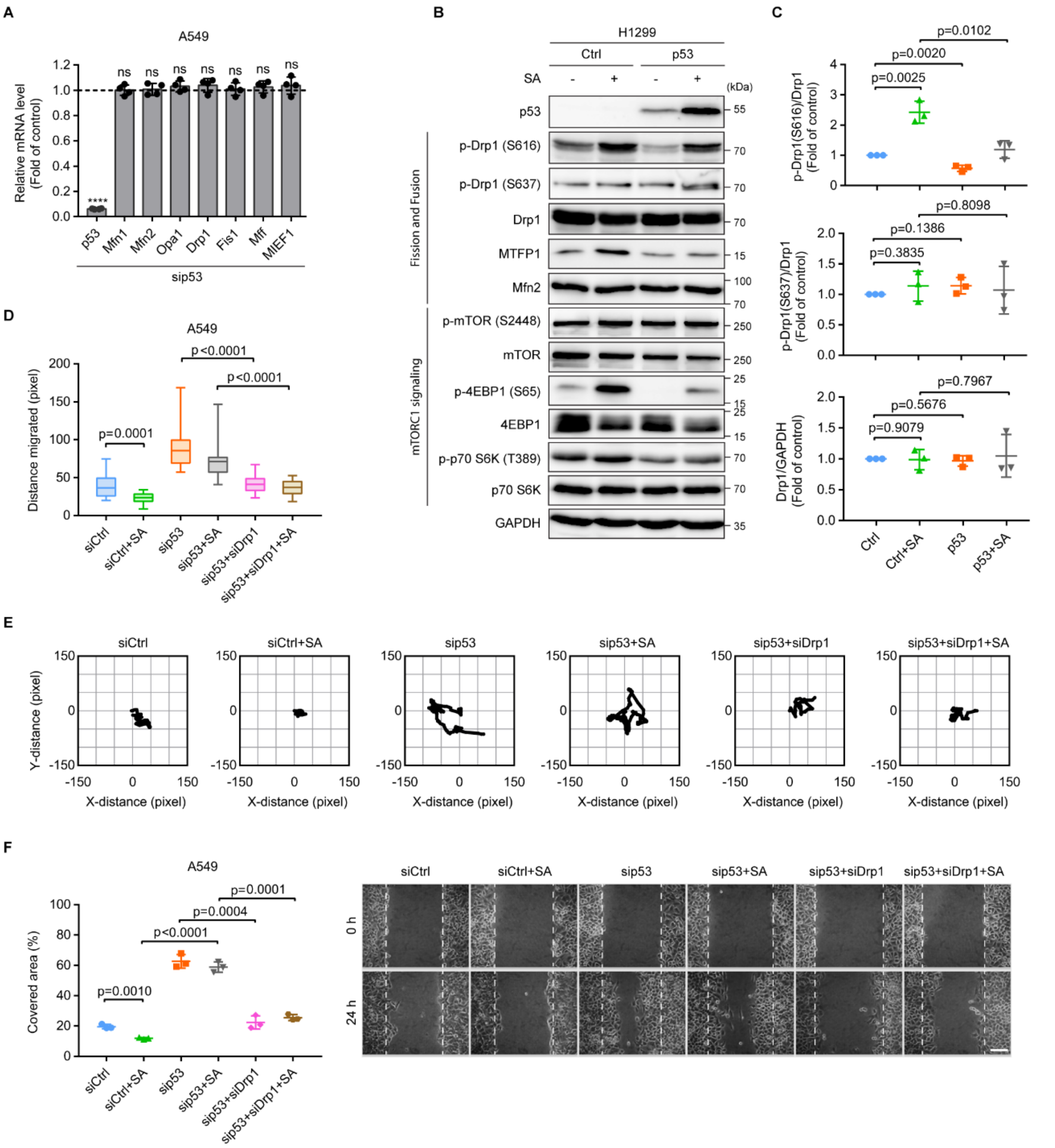


**Fig. S2** p53 suppresses cell motility by inhibiting the pro-fission phosphorylation of Drp1. **A** qRT-PCR analysis of the mRNA levels of p53 and genes involved in mitochondrial fusion (Mfn1, Mfn2, and Opa1) and fission (Drp1, Fis1, Mff, and MIEF1) in sip53 A549 cells. Results are expressed relative to those in siCtrl A549 cells (dashed line). **B** Immunoblot of the indicated proteins in Ctrl and p53 H1299 cells with and without 20 μM SA treatment for 24 h. GAPDH was used as a loading control. **C** Quantification of the levels of Drp1, p-

Drp1 (S637), and p-Drp1 (S616) in **B**. **D**, **E** Migration distance (**D**) and representative trajectories (**E**) of siCtrl (n = 29), siCtrl+SA (n = 17), sip53 (n = 13), sip53+SA (n = 15), sip53+siDrp1 (n = 24), and sip53+siDrp1+SA (n = 24) A549 cells. **F** Quantification (left) and representative images (right) of the area in a wound-healing assay covered by siCtrl, siCtrl+SA, sip53, sip53+SA, sip53+siDrp1, and sip53+siDrp1+SA A549 cells. Scale bar: 100 μm. Error bars represent mean ± SD. Data were analyzed by two-tailed unpaired Student's t test. ****, $p < 0.0001$; ns, not significant.

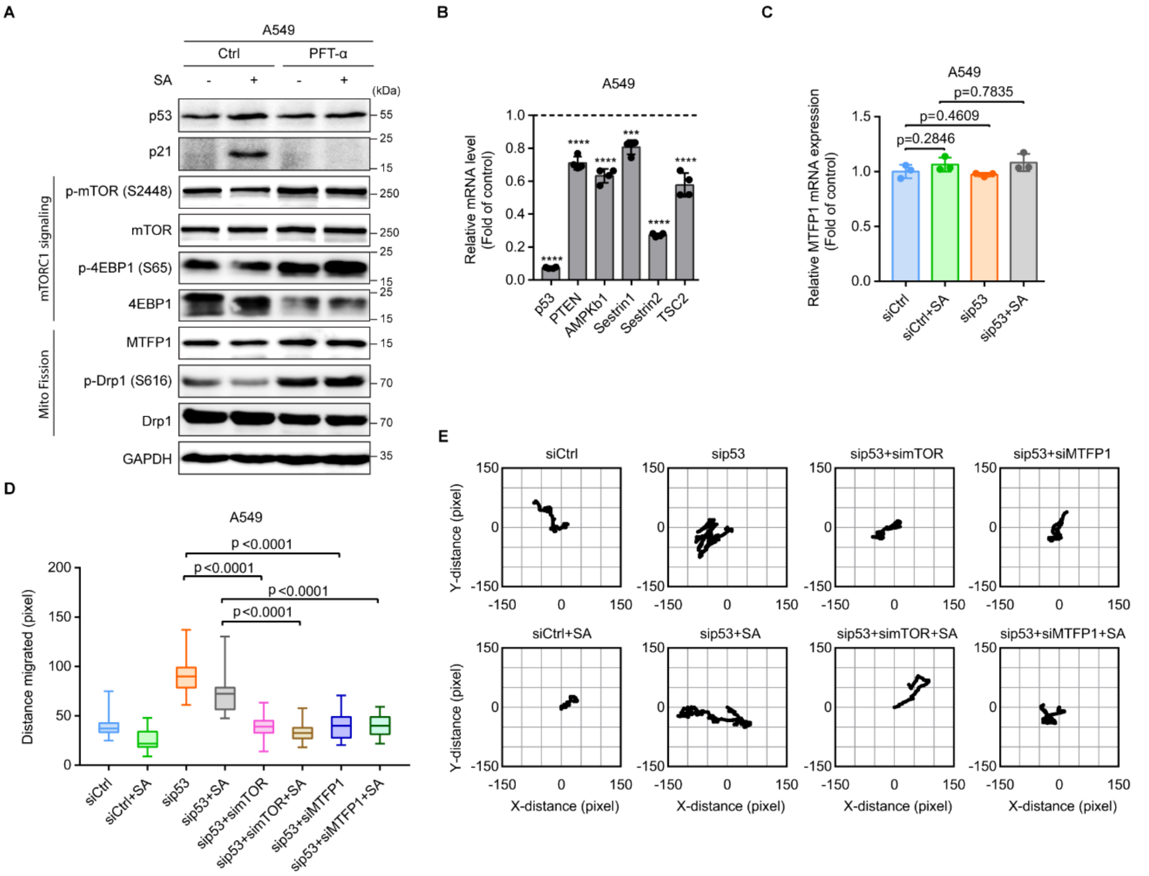


**Fig. S3** p53 transcriptional activity participates in regulating mTORC1-controlled MTFP1 protein levels affecting cell motility. **A** Immunoblot of the indicated proteins in control (Ctrl) and PFT-α-treated (PFT-α) A549 cells with and without 20 μM SA treatment for 24 h. GAPDH was used as a loading control. **B** qRT-PCR analysis of the mRNA levels of p53 and p53 downstream target genes in sip53 A549 cells. Results were expressed relative to those in siCtrl A549 cells (dashed line). **C** qRT-PCR analysis of mRNA expression of MTFP1 in siCtrl and sip53 A549 cells with and without 20 μM SA treatment for 24 h. **D**, **E** Migration distance (**D**) and representative trajectories (**E**) of siCtrl (n = 28), siCtrl+SA (n = 17), sip53 (n = 13), sip53+SA (n = 14), sip53+simTOR (n = 33), sip53+simTOR+SA (n = 28), sip53+siMTFP1 (n = 32), and sip53+siMTFP1+SA (n = 20) A549 cells. Error bars represent mean ± SD. Data were analyzed by two-tailed unpaired Student’s t test. ***, $p < 0.001$; ****, $p < 0.0001$.

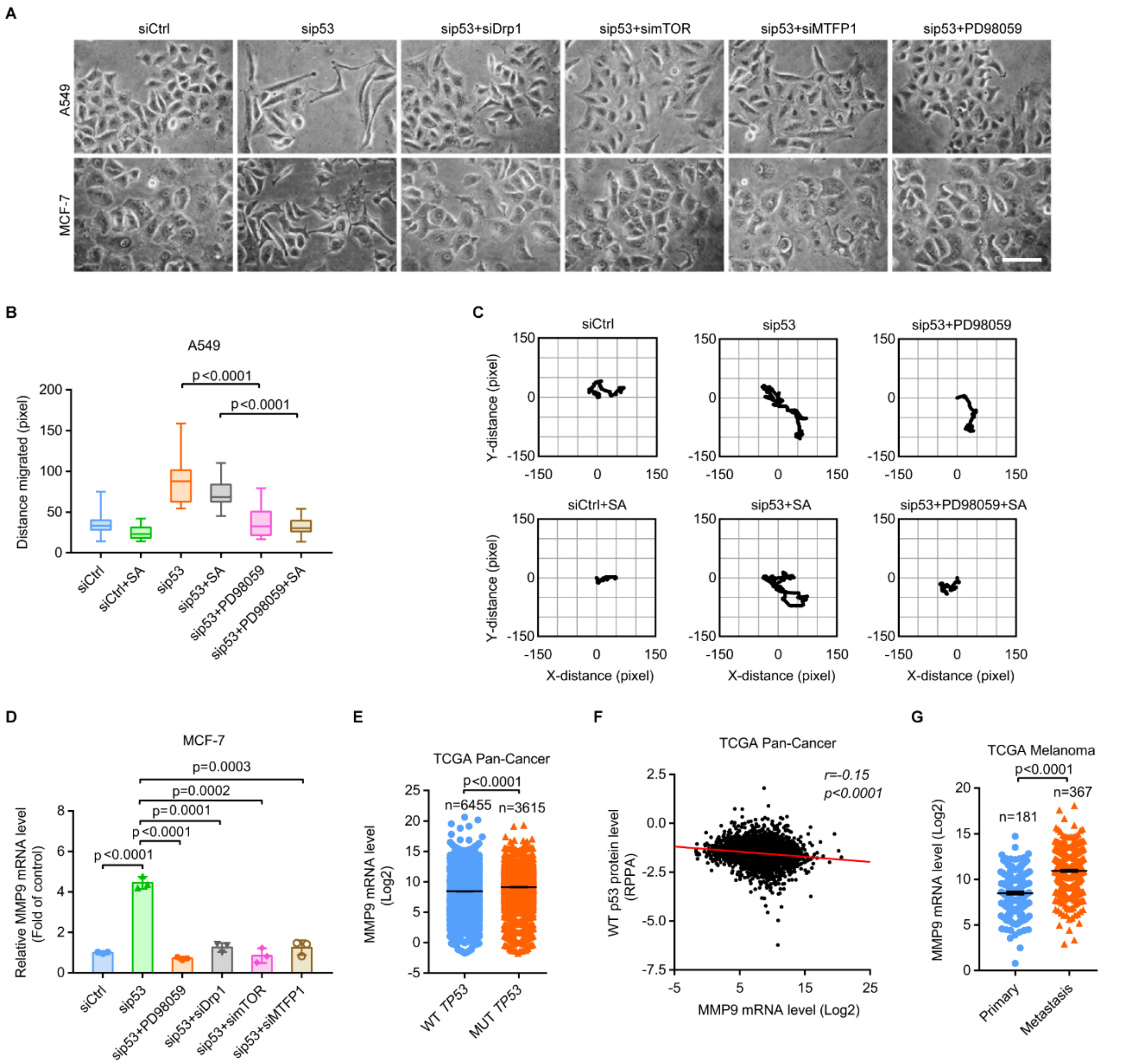


**Fig. S4** p53 controls cell motility and MMP9 expression through mTOR/MTFP1/Drp1/ERK1/2 signaling axis. **A** Phase-contrast imaging of siCtrl, sip53, sip53+siDrp1, sip53+simTOR, sip53+siMTFP1, and sip53+PD98059 A549 and MCF-7 cells. Scale bar: 100 μm. **B**, **C** Migration distance (**B**) and representative trajectories (**C**) of siCtrl (n = 31), siCtrl+SA (n = 19), sip53 (n = 13), sip53+SA (n = 14), sip53+ PD98059 (n = 20), and sip53+PD98059+SA (n = 35) A549 cells. **D** qRT-PCR analysis of MMP9 mRNA expression in siCtrl, sip53, sip53+PD98059, sip53+siDrp1, sip53+simTOR, and sip53+siMTFP1 MCF-7 cells. **E** MMP9 mRNA expression in tumors having WT and MUT

*TP53*. **F** Correlation between p53 protein levels and MMP9 mRNA levels in tumors having WT *TP53* (n = 4 538 samples). **G** MMP9 mRNA levels in primary and distant metastatic melanoma. Data were extracted from TCGA (**E**-**G**). Error bars represent mean ± SD (**D**) or SEM (**E**, **G**). Data were analyzed by two-tailed unpaired Student's t test.

### Supplementary Tables

#### Table S1. siRNA sequences

| Target protein | Oligo ID | Sequence (5'-3') |
|---|---|---|
| p53 | VHS40367 | Sense: CCAGUGGUAAUCUACUGGGACGGAA |
| | | Antisense: UUCCGUCCCAGUAGAUUACCACUGG |
| p53 | VHS40366 | Sense: CCAUCCACUACAACUACAUGUGUAA |
| | | Antisense: UUACACAUGUAGUUGUAGUGGAUGG |
| Drp1 | HSS115288 | Sense: CCUGCUUUAUUUGUGCCUGAGGUUU |
| | | Antisense: AAACCUCAGGCACAAAUAAAGCAGG |
| mTOR | HSS103825 | Sense: AGGACGCUCACAUUGCUAGAUGUGG |
| | | Antisense: CCACAUCUAGCAAUGUGAGCGUCCU |
| MTFP1 | HSS182106 | Sense: GGGAUACCUGGGCUAUGCCAAUGAG |
| | | Antisense: CUCAUUGGCAUAGCCCAGGUAUCGC |

**Table S2. List of primers used for qRT-PCR**

| Target protein | Sequence (5’-3’) |
|---|---|
| p53 | Forward: AAGGAAATTTGCGTGTGGAGT |
| | Reverse: AAAGCTGTTCCGTCCCAGTA |
| Mfn1 | Forward: GAGGTGCTATCTCGGAGACAC |
| | Reverse: GCCAATCCCACTAGGGAGAAC |
| Mfn2 | Forward: CACATGGAGCGTTGTACCAG |
| | Reverse: TTGAGCACCTCCTTAGCAGAC |
| Opa1 | Forward: TGTGAGGTCTGCCAGTCTTTA |
| | Reverse: TGTCCTTAATTGGGGTCGTTG |
| Drp1 | Forward: ACCCGGAGACCTCTCATTCT |
| | Reverse: TGACAACGTTGGGTGAAAAA |
| Fis1 | Forward: GATGACATCCGTAAAGGCATCG |
| | Reverse: AGAAGACGTAATCCCGCTGTT |
| Mff | Forward: CACCACCTCGTGTACTTACGC |
| | Reverse: GTCTGCCAACTGCTCGGATTT |
| MIEF1 | Forward: CACGGCCATTGACTTTGTGC |
| | Reverse: TCGTACATCCGCTTAACTGCC |
| PTEN | Forward: AGTTCCCTCAGCCGTTACCT |
| | Reverse: AGGTTTCCTCTGGTCCTGGT |
| AMPKβ1 | Forward: TCCGATGTGTCTGAGCTGTC |
| | Reverse: GTTCAGCATGACGTGATTGG |
| Sestrin1 | Forward: AGCCCATAGACCTTGGCTTA |
| | Reverse: TCCACACTGTGATTGCCATT |

| | |
|---|---|
| Sestrin2 | Forward: TGCTGTGCTTTGTGGAAGAC |
| | Reverse: GCTGCCTGGAACTTCTCATC |
| TSC2 | Forward: TGCAAGCCGTCTTCCACAT |
| | Reverse: ATGGACACAAAGTCGTTGC |
| MTFP1 | Forward: CCATCCCCATCATTATCCAC |
| | Reverse: TTCCCCACTGTTGGGTAGAG |
| GAPDH | Forward: TGCACCACCAACTGCTTAGC |
| | Reverse: GGCATGGACTGTGGTCATGAG |
